# NK-RCPP: a receptor-conditioned computational workflow for prioritizing neurokinin-receptor peptide candidates

**DOI:** 10.64898/2026.10.23.761317

**Authors:** Xiaosong Zhou, Haohong Zhang, Kang Ning

**Author notes:** These authors contributed equally (Xiaosong Zhou, Haohong Zhang). Correspondence should be addressed to K. N.

## Abstract

Prioritizing peptides for closely related G protein-coupled receptors requires comparing target-directed scores with evidence from other structural contexts. We applied the Neurokinin Receptor-Conditioned Peptide Prioritization workflow (NK-RCPP) to 96 generated candidates for neurokinin receptors NK1R, NK2R and NK3R. The retrospective analysis combined receptor-matched on-target REF15 scores with 4,518 paired structural-neighbour comparisons covering 82 candidates. Lower on-target scores than the matched controls were observed for 14/32 NK1R, 23/32 NK2R and 6/32 NK3R candidates. Adding the neighbour-score criterion retained 16 of the 33 evaluable on-target-favoured candidates, reducing that pool by 51.5%. At matched receptor-specific shortlist sizes, half of the candidate identities differed from on-target-only selection. NK2R-04 and NK2R-19 remained selected at the higher neighbour-fraction threshold of 0.75. Fourteen NK2R candidates lacked neighbour summaries, leaving their joint-rule status unresolved. Sequence-frequency and embedding analyses described the candidates within a 1,048-peptide comparison space, and an exploratory 10-peptide NK2R comparison showed differences in standardized scores with uncertain score-potency agreement. These findings show how paired structural-neighbour scoring and threshold choice shape candidate selection within a receptor family. The resulting shortlist provides a defined set for testing whether joint prioritization enriches for active, receptor-preferred peptides.

**Highlights:**

- NK-RCPP combines receptor-matched scores and paired structural-neighbour comparisons to prioritize peptides across three neurokinin receptors.
- On-target REF15 scores were lower than the matched controls for 14/32 NK1R, 23/32 NK2R and 6/32 NK3R candidates.
- Joint prioritization retained 16/33 evaluable on-target-favoured candidates and changed half of the selections at matched shortlist sizes.
- NK2R-04 and NK2R-19 remained selected at the higher neighbour-fraction threshold; missing summaries left 14 NK2R candidates unclassified.

## Introduction

G protein-coupled receptors (GPCRs) constitute the largest family of targets for approved medicines, and peptide-responsive GPCRs translate extracellular signals into physiological responses [1]. The tachykinin system illustrates the challenge of designing ligands for closely related receptors. Substance P, neurokinin A (NKA) and neurokinin B (NKB) preferentially activate NK1R, NK2R and NK3R, respectively, and regulate nociception, inflammation, smooth-muscle activity and neuroendocrine signalling [2]. The clinical efficacy of NK1R blockade in chemotherapy-induced nausea and vomiting and NK3R blockade in menopausal vasomotor symptoms establishes the pharmacological relevance of this receptor family [3, 4]. Prioritizing new peptide candidates requires a way to compare receptor-directed scores while accounting for the shared recognition features of the three subtypes.

Subtype discrimination depends on conserved tachykinin recognition and receptor-specific contacts. Tachykinins share the amidated C-terminal pharmacophore Phe-X-Gly-Leu-Met-NH2, whereas receptor preference also involves the divergent peptide N terminus, extracellular loops and receptor N-terminal region. Agonist-bound structures of NK1R and NK3R and structure-function analysis of NK1R and NK2R reveal a conserved recognition core with subtype-specific interaction networks [5–7]. Experiments with modified NKA peptides further distinguish affinity, potency, efficacy and subtype selectivity as separate properties [8]. These findings motivate receptor-matched comparisons during computational prioritization and receptor-family measurements during experimental characterization. Sequence generation and structural modelling address different parts of peptide discovery. Peptides provide chemical and conformational diversity, while their flexibility, proteolytic instability and context-dependent pharmacology complicate screening [9]. Protein language models generate functional protein sequences and target-conditioned peptide proposals [10, 11]. PepMLM conditions peptide generation on the target sequence [10], and structure-based methods sample peptide-receptor conformations and rank them with model scores [12–15]. The interpretation of these scores depends on receptor preparation, peptide length and conformational sampling. A benchmark of 113 known GPCR-peptide and GPCR-protein complexes found confidence overestimation for misplaced ligands in several structure-prediction methods [16]. Candidate selection therefore benefits from examining how rankings change across scoring contexts.

For a receptor-family screen, the practical question is how comparative scores affect the candidates selected for experiments. Ranking solely by the target score can favour peptides that also receive lower scores on other structures. Paired candidate-control comparisons across structural neighbours offer a second ranking dimension. Its contribution can be assessed by comparing the resulting shortlist with on-target-only selection at the same testing budget, then examining sensitivity to thresholds and missing comparisons. This makes the consequences of each selection rule explicit. Generative design has expanded functional sequence space through enzyme generation with ProteinGAN [17] and ProGen [11], inverse folding with ProteinMPNN [18] and programmable protein sampling with Chroma [19]. Structure-guided approaches have produced protein binders from target structures [20], including designs generated with RFdiffusion [21] and proteins that bind bioactive helical peptides [22]. BindCraft integrates design and filtering to produce functional protein binders [23]. For peptide generation, PepPrCLIP uses contrastive language modelling [24], whereas PepMLM uses target-conditioned masked language modelling [10]. These studies establish generation-and-selection workflows [10, 21, 23]. Here, we examine how paired structural-neighbour scores alter candidate selection in a neurokinin-receptor screen.

We used NK-RCPP to compare 96 receptor-conditioned peptide candidates with their receptor-matched ligand controls and to evaluate how structural-neighbour scoring changes the shortlist (**Fig. 1**). The analysis combined on-target scores with paired neighbour comparisons, quantified candidate replacement at matched receptor-specific shortlist sizes, and examined threshold sensitivity and missing NK2R summaries. Position-wise sequence frequencies, an exploratory NK2R peptide comparison and an embedding projection provided additional context for the selected candidates. The study links each priority to its score criteria and defines the candidate sets for subsequent experimental comparison.

**Fig. 1.**
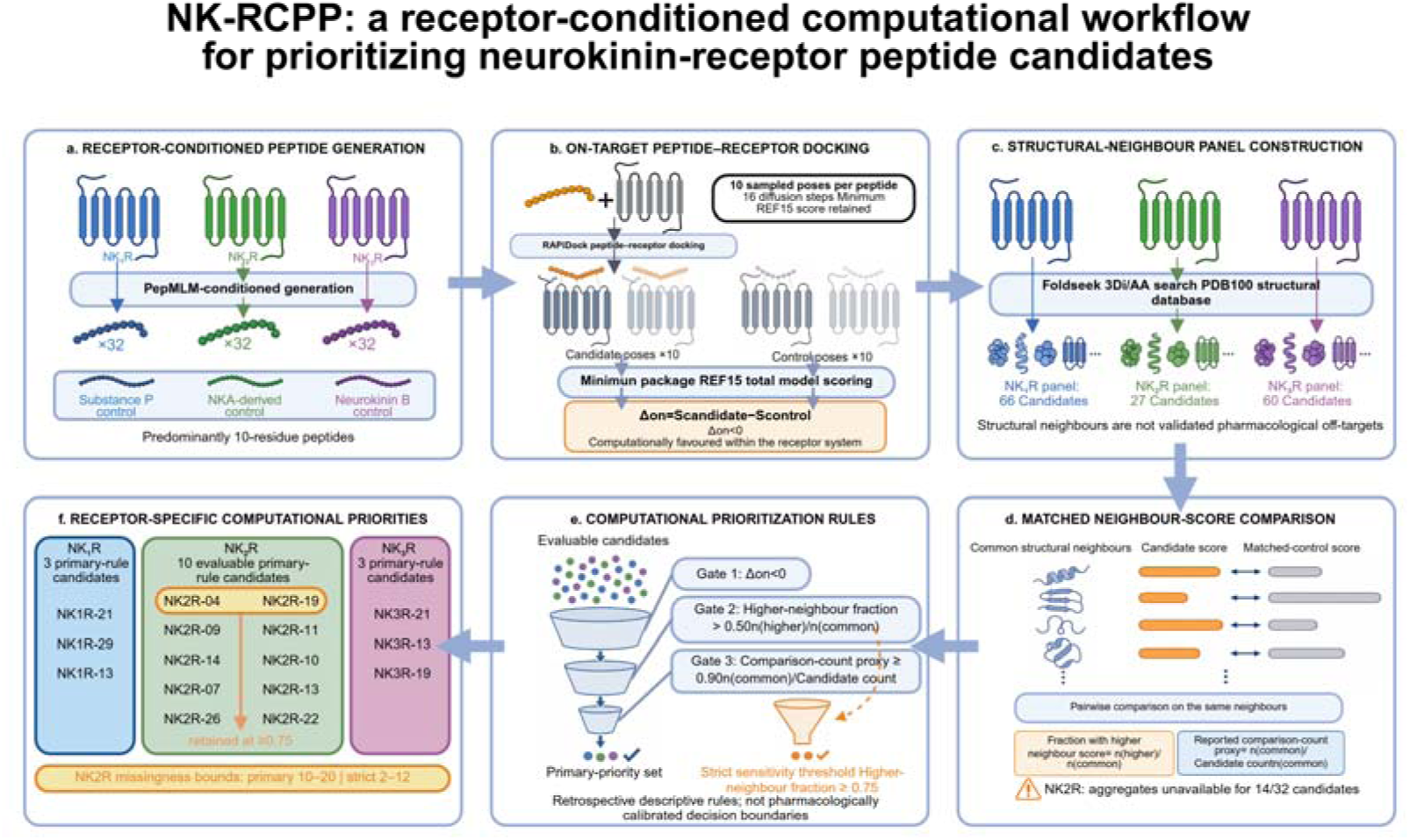
Neurokinin Receptor–Conditioned Peptide Prioritization (NK-RCPP) workflow. a, PepMLM generated 32 candidates per receptor, predominantly ten-residue peptides. Receptor-matched controls were Substance P, an NKA-derived peptide and neurokinin B. b, RAPiDock sampled peptide-receptor poses; the minimum REF15 total model score was retained. Delta-on is the candidate score minus the matched control score. c, Foldseek retrieval and redundancy reduction defined structural-neighbour panels with 66 NK1R, 27 NK2R and 60 NK3R entries (Table S3). d, Paired candidate-control scores yielded n_common, n_higher, the neighbour-score fraction and comparison-count proxy. Fourteen NK2R candidates lacked neighbour summaries. e, The primary rule required Delta-on < 0 and fraction > 0.50, with count proxy ≥0.90 as a completeness check; all 82 evaluable candidates had proxy 1.00. The higher-threshold rule required fraction ≥0.75. f, The primary rule retained three NK1R, ten NK2R and three NK3R candidates; NK2R-04 and NK2R-19 also passed the higher threshold. Missing-outcome bounds for NK2R were 10–20 primary-rule and 2–12 higher-threshold candidates.

## Results

### On-target scoring identified candidates within each receptor set

The NK-RCPP dataset contained 32 PepMLM-generated candidates and one receptor-matched ligand control for each of NK1R, NK2R and NK3R (Fig. 1). We used the minimum of ten on-target REF15 total model scores per complex and calculated Delta-on as the candidate score minus the matched control score. Joint prioritization combined Delta-on < 0 with a neighbour-score fraction > 0.50, defined as the proportion of paired structural-neighbour comparisons in which the candidate score exceeded the control score. A comparison-count proxy of at least 0.90 checked panel completeness.

On-target REF15 scores were lower than the matched control for 23/32 NK2R candidates (71.9%), 14/32 NK1R candidates (43.8%) and 6/32 NK3R candidates (18.8%) (**Fig. 2a** and **Supplementary Figs. S1 and S8**). The lowest scores within each receptor set occurred for NK1R-18 (517.29 versus 574.39 for Substance P; Delta-on = -57.10), NK2R-27 (636.35 versus 771.13 for the NKA-derived control; Delta-on = -134.78) and NK3R-21 (1965.25 versus 2240.54 for NKB; Delta-on = -275.29). Comparisons were made within each receptor system; the NK1R set paired ten-residue candidates with eleven-residue Substance P.

**Fig. 2.**
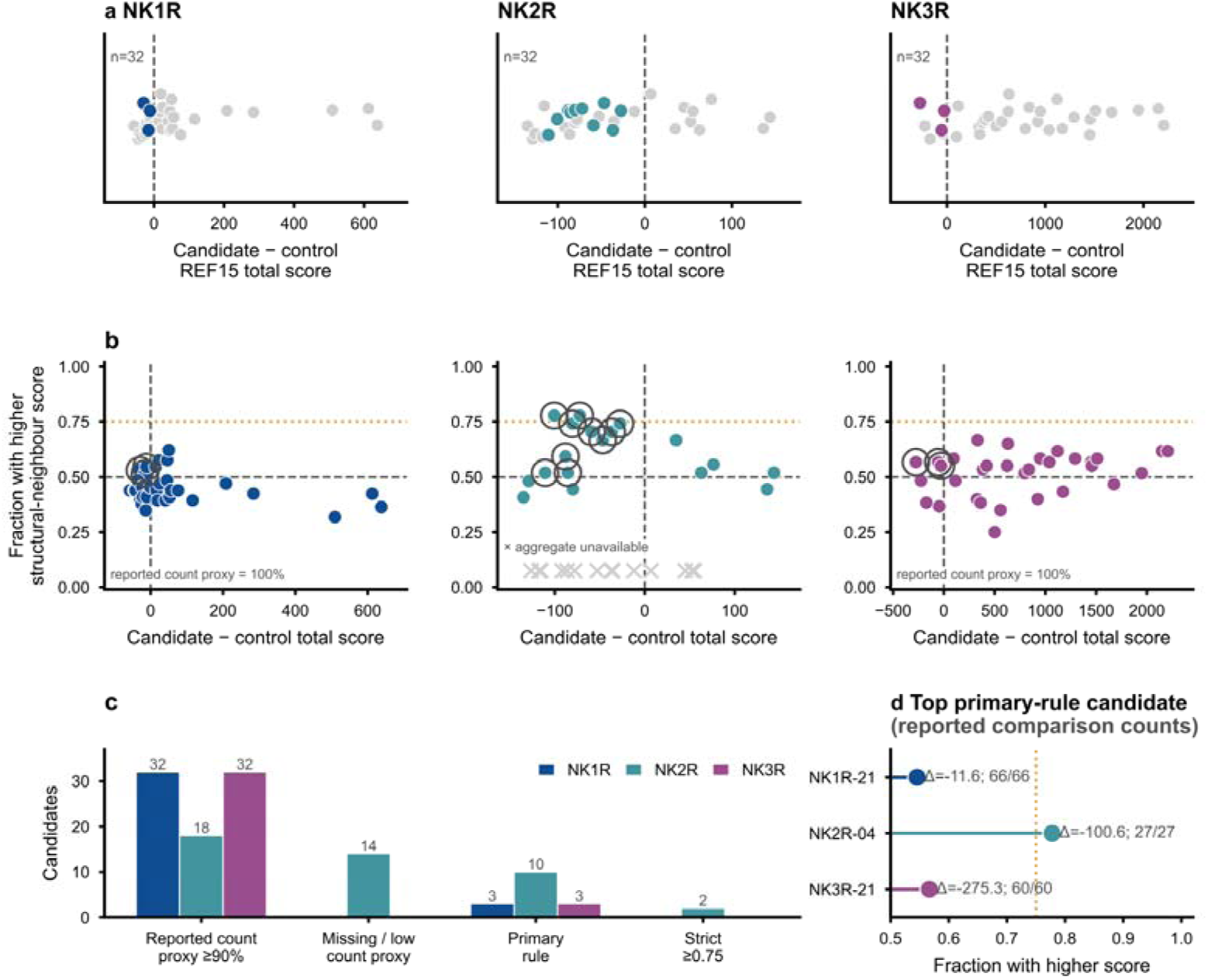
Receptor-specific on-target scores and joint prioritization. a, Candidate-minus-control minimum REF15 total model scores for 32 candidates per receptor; values below zero denote a lower candidate score within the receptor system. b, Delta-on versus the neighbour-score fraction. Dashed lines mark Delta-on = 0 and fraction = 0.50; the dotted amber line marks 0.75. Filled receptor-coloured points have a comparison-count proxy ≥0.90, and outlines identify primary-rule candidates. Crosses show NK2R candidates without neighbour summaries. c, Counts meeting the completeness check, missing or below the proxy threshold, passing the primary rule and passing the ≥0.75 rule. d, Top primary-rule candidate per receptor, ordered by fraction and Delta-on; labels report Delta-on and n_common divided by the Table S3 panel size.

Structural-neighbour summaries were available for 32/32 NK1R, 18/32 NK2R and 32/32 NK3R candidates, covering 82 candidates and 4,518 paired candidate-control comparisons (**Fig. 2b** and **Supplementary Figs. S1-S3**). Each evaluable candidate had results for its complete receptor-associated panel of 66, 27 or 60 structures, respectively. The comparison-count proxy was therefore 1.00 for all 82 candidates, and the ≥0.90 completeness check excluded no additional candidate.

### Joint prioritization changed candidate identities at matched shortlist sizes

The primary joint rule retained three NK1R, ten NK2R and three NK3R candidates (**Fig. 3a-c** and **Supplementary Figs. S2 and S5**). The set comprised NK1R-21, NK1R-29 and NK1R-13; NK2R-04, NK2R-19, NK2R-14, NK2R-07, NK2R-26, NK2R-09, NK2R-11, NK2R-10, NK2R-13 and NK2R-22; and NK3R-21, NK3R-13 and NK3R-19. The retained NK1R neighbour-score fractions were 36/66, 35/66 and 34/66. NK2R-04 and NK2R-19 each had a higher score than the control in 21/27 neighbour comparisons (0.778), with Delta-on values of -100.60 and -72.21. The retained NK3R fractions were 34/60, 34/60 and 33/60, with Delta-on values of -275.29, -55.26 and -28.47.

**Fig. 3.**
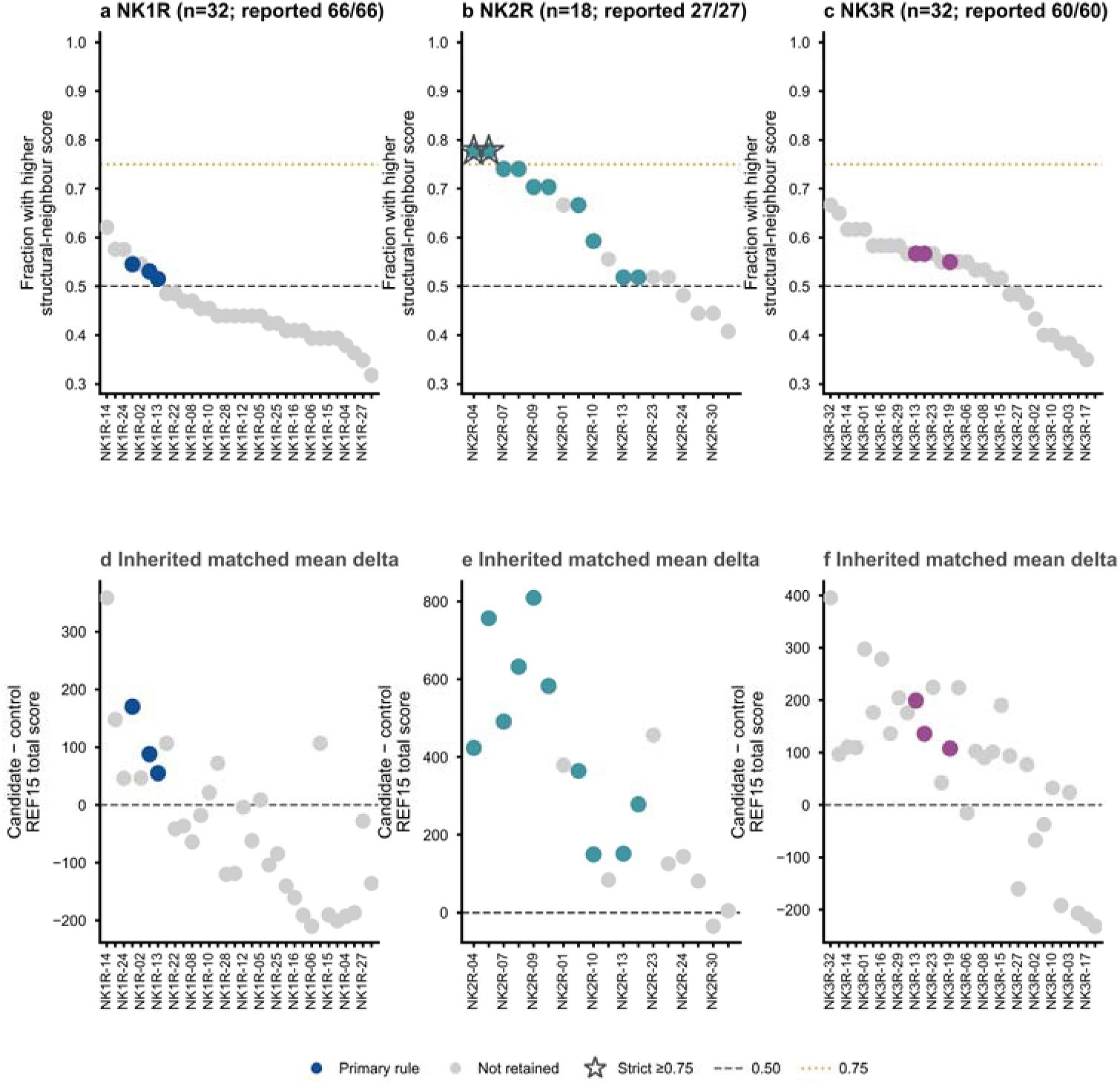
Candidate rankings by structural-neighbour score fraction and paired differences. a–c, Evaluable NK1R, NK2R and NK3R candidates ranked by the fraction of structural-neighbour comparisons with a higher candidate REF15 total model score than the matched control. Receptor-coloured points satisfy the primary joint rule, grey points fall outside it, and stars identify candidates passing the ≥0.75 sensitivity rule. Dashed and dotted lines mark fractions of 0.50 and 0.75. Titles give the comparison count relative to the receptor-specific Table S3 panel size. d–f, Mean paired candidate-minus-control score differences for the same candidates. Median differences are shown in **Supplementary Fig. S2**; for NK1R-21, the mean and median were 170.04 and 7.20, respectively.

Of the 43 candidates with lower on-target scores than their controls, 33 had neighbour summaries and ten remained unclassified under the joint rule. Retaining 16/33 evaluable candidates reduced the on-target-favoured pool by 51.5%. At equal shortlist sizes of three NK1R, ten NK2R and three NK3R candidates, the joint and on-target-only selections shared 0/3, 7/10 and 1/3 candidates, respectively (8/16 overall). Thus, adding the neighbour criterion replaced half of the candidate identities at a fixed testing budget.

The two score summaries favoured different leading candidates (**Fig. 3a-f**). NK1R-18 had the lowest NK1R on-target score but a neighbour-score fraction of 29/66 (0.439); NK2R-27 had the lowest NK2R on-target score but a fraction of 11/27 (0.407). Conversely, NK1R-14 had the highest qualified NK1R fraction, 41/66 (0.621), while scoring 50.23 units above its matched control on target. The first two candidates failed the neighbour criterion, whereas NK1R-14 failed the on-target criterion. Representative pocket renderings show the predicted candidate and control conformations (**Fig. 5a-d** and **Supplementary Figs. S4, S5f, S6f and S8d**). Mean and median paired differences also captured different aspects of the score distributions: for NK1R-21, the mean was 170.04 and the median 7.20, with individual differences from −5,858.24 to 7,639.93. Its fraction of 36/66 summarizes the direction of those differences independently of their magnitude (**Supplementary Figs. S2 and S5**).

### Thresholds and missing comparisons affected shortlist size

Candidate counts depended on both the neighbour-fraction threshold and missing NK2R comparisons (Fig. 2c,d). Raising the cutoff from >0.50 to >0.55 removed all NK1R candidates; requiring ≥0.75 retained only NK2R-04 and NK2R-19. Neighbour summaries were missing for 14 NK2R candidates, including ten with Delta-on < 0. Assigning none or all of these ten candidates a passing neighbour fraction gives NK2R count bounds of 10–20 under the primary rule and 2–12 under the higher-threshold rule. These ranges describe the possible counts given the missing comparisons.

### Generated peptides showed receptor-stratified sequence patterns

The exact ten-residue candidate sets had distinct position-wise amino-acid distributions across the three receptors (**Fig. 4** and **Supplementary Fig. S9**). At a per-position frequency threshold of 0.30, the consensus strings were SVPRQQCFGF for NK1R, PKPRQCCFQR for NK2R and DTPYQQSNGF for NK3R. NK2R-02 (RRVPRQCCFQR) contained eleven residues and was excluded from the position-wise heatmap, leaving 31 NK2R candidates. It remained in raw-score summaries but was excluded from length-standardized analyses because no eleven-residue random-control baseline was available. These frequency summaries describe sequence patterns within the generated sets.

**Fig. 4.**
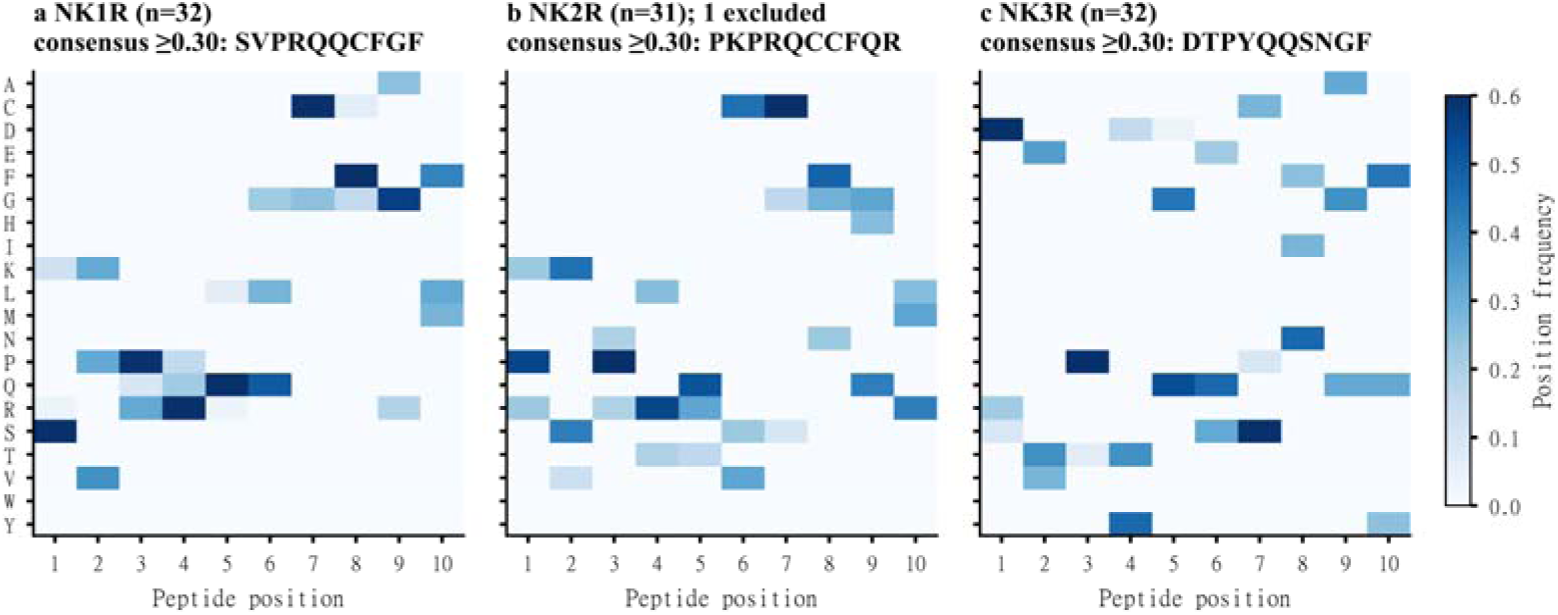
Position-wise amino-acid frequencies in generated peptides. a–c, Frequencies of the 20 standard amino acids at each position of exact ten-residue NK1R, NK2R and NK3R candidates. Heatmaps share a frequency scale of 0–0.60 and the amino-acid order ACDEFGHIKLMNPQRSTVWY. A consensus character is shown when the most frequent residue reaches at least 0.30, yielding SVPRQQCFGF, PKPRQCCFQR and DTPYQQSNGF, respectively. All 32 NK1R and 32 NK3R candidates were included. The eleven-residue NK2R-02 was excluded, leaving 31 NK2R candidates.

### Exploratory NK2R comparisons showed differences in score distributions

The exploratory NK2R comparison included ten competition peptides spanning six, nine and fifteen residues (**Supplementary Figs. S6 and S7**). Nine of ten had a length-stratified z-score below zero, 5/10 had Delta-on < 0, 8/10 had a neighbour-score fraction > 0.50, and 3/10 met the joint score rule. Among the 18 evaluable study NK2R candidates, 14 passed the fraction criterion and ten passed the joint rule; 3/5 competition nine-mers passed the joint rule. With competition peptides as the first group, raw-score comparisons gave U = 202, rank-biserial r = +0.263 and P = 0.2204 (10 versus 32), and fraction comparisons gave U = 119, r = +0.322 and P = 0.1699 (10 versus 18, with ties). Excluding the eleven-residue NK2R-02 from standardization, competition z-scores were lower than those of the 31 study ten-mers (medians −0.92 versus 0.48; U = 40, r = −0.742, P = 5.09 × 10□□), including 270/310 cross-group score pairs (87.1%). The estimated correlation with workbook-reported potency was uncertain (Spearman rho = +0.42, P = 0.23, n = 10). These exploratory comparisons describe groups with different lengths, residue compositions and sampling frames.

### Sequence projection placed candidates among comparison peptides

The ESM-2 (Evolutionary Scale Modeling 2) projection placed the study candidates within a comparison set of 1,048 peptides (**Fig. 5e-h**). Representations were normalized to unit length and projected into two dimensions with UMAP (Uniform Manifold Approximation and Projection). A separated island contained 92 sequences of 7-10 residues, while the main projection contained 956 sequences of 3-40 residues; 4/10 competition peptides fell in the island. In the plotted coordinates, the median distance to the nearest retained leaderboard peptide was 0.04 for the ten primary-rule NK2R candidates and 0.07 for all 99 study entries, including the three controls. This projection describes the sequence context of the candidates alongside the position-wise frequency summaries.

**Fig. 5.**
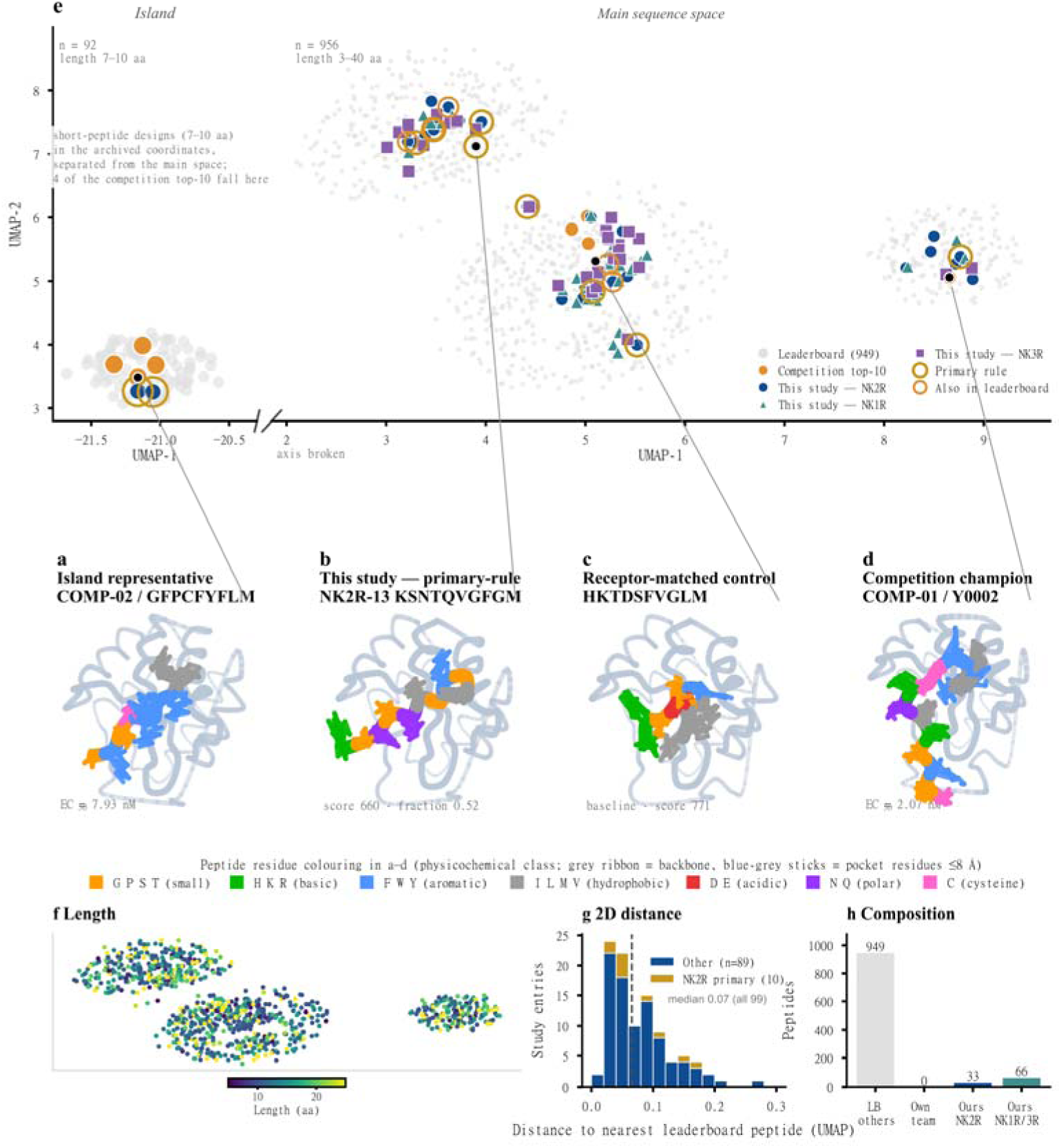
Peptide-pocket renderings and sequence-space context. a–d, Computational peptide-pocket renderings for COMP-02, the primary-rule candidate NK2R-13, the receptor-matched NK2R control and COMP-01. Residue colours denote physicochemical classes. e, UMAP projection of L2-normalized ESM-2 embeddings, with a broken UMAP-1 axis separating the 92-sequence island from the 956-sequence main projection. f, Peptide lengths across the main projection. g, Two-dimensional distance to the nearest retained leaderboard peptide for all 99 study entries, including controls, and for the ten primary-rule NK2R candidates; median distances were 0.07 and 0.04, respectively. h, Composition of the plotted peptide groups.

## Discussion

NK-RCPP shows how paired structural-neighbour scoring changes peptide priorities within the neurokinin receptor family. The joint rule retained 16 of 33 evaluable candidates favoured by on-target scoring and replaced half of the candidate identities when shortlist sizes were matched. These changes identify a practical consequence of combining score summaries: the choice of scoring context affects which peptides enter an experimental portfolio, even at a fixed testing budget.

The neurokinin family provides a biologically relevant setting for this comparison. Clinical applications of NK1R and NK3R antagonists establish the importance of these receptors [3, 4], while their shared tachykinin recognition features motivate receptor-matched peptide comparisons. The generated sets displayed different position-wise sequence patterns, giving a sequence-level description of the candidate pools. These patterns complement the score-based shortlist by showing which residues recur within each receptor-conditioned set.

Disagreement between the score summaries explains the changes in candidate selection. NK1R-18 and NK2R-27 led their on-target rankings but fell below the neighbour-fraction threshold, whereas NK1R-14 led the qualified NK1R neighbour ranking but failed the on-target rule. The paired neighbour comparisons thus expose score patterns that an on-target ranking alone would leave out of the selection decision. Target-conditioned generation and multistage design-and-filter workflows are established capabilities [10, 21, 23]; this study quantifies their selection consequences in a receptor-family dataset through matched controls, equal-size shortlist comparisons and threshold sensitivity.

The exploratory NK2R comparison and sequence projection provide complementary context for the shortlist. Competition peptides had lower standardized scores than the study ten-mers, with an uncertain score-potency correlation. Each length baseline contained five random peptides, residue pools differed between lengths, and the ten-residue pool was based on NK2R-01 and the NKA-derived control. These sampling choices shape the standardized comparison. ESM-2 and UMAP place the candidates among reference sequences [25, 26], while the reported distances describe the two-dimensional projection. Neither analysis contributed a criterion to the joint selection rule.

Threshold sensitivity identifies priorities that depend on small changes in the decision rule. Raising the neighbour-score fraction cutoff from >0.50 to >0.55 removed all NK1R candidates, whereas NK2R-04 and NK2R-19 remained selected at ≥0.75. Candidates close to a cutoff therefore warrant comparison with those on the other side of the boundary. Missing neighbour summaries for 14 NK2R candidates also leave part of the on-target-favoured pool unresolved; the reported count bounds make this dependence explicit.

The findings concern one fixed dataset evaluated with retrospective thresholds and REF15 total model scores. These scores describe modelled complexes, and the structural-neighbour panels contain receptor-related structures, unrelated enzymes and engineered chimeras; they do not directly measure affinity, receptor selectivity or safety. Receptor constructs and states differ, including the NK1R glycogen-synthase fusion and its comparison of ten-residue candidates with an eleven-residue control. Raw pose coordinates, chemical-preparation records, historical software versions and Foldseek filtering logs were unavailable, as were NK2R pseudo-perplexity values and experimental activity labels for the 96 candidates. These features define the scope of the present computational comparison [12, 15, 16].

A direct next test is to compare joint-rule, on-target-only and native PepMLM-ranked selections at matched receptor-specific testing budgets, including the two NK2R candidates retained at the higher threshold and candidates near the decision boundaries. Binding measurements and concentration-response assays across NK1R, NK2R and NK3R in matched cellular backgrounds would test whether the altered shortlist enriches for active, receptor-preferred peptides. Functional measurements can then distinguish potency, efficacy and signalling bias [8, 27].

## Conclusion

NK-RCPP links receptor-matched target scores with paired structural-neighbour comparisons to define peptide priorities across the neurokinin receptor family. Joint selection changed half of the candidate identities at matched shortlist sizes, with NK2R-04 and NK2R-19 retained under the higher neighbour-fraction threshold. Threshold sensitivity and missing-outcome bounds specify how the shortlist depends on the available evidence. The resulting candidate sets support a direct experimental comparison of joint and single-ranking selection strategies.

## Methods

### Study design

We performed a retrospective descriptive analysis of a fixed computational screening dataset. NK-RCPP comprises receptor-conditioned generation, on-target docking, structural-neighbour comparison and deterministic prioritization. The analysis used candidate tables, score summaries, panel annotations and workflow scripts, together with per-complex scores from the public repository at commit 4a04cab13476e30ba5cd1a24ea876ca6d9305998. Scores were joined by receptor, peptide sequence and neighbour identity to reconstruct 4,518 paired comparisons, 82 candidate aggregates and their panel membership. We also checked the ten on-target score columns for each study complex. Generation, docking, structure prediction, Foldseek searches and embedding generation were not rerun. The receptor-specific analysis was descriptive; exploratory inferential tests were confined to the external NK2R comparison. The primary (>0.50) and higher (≥0.75) neighbour-fraction thresholds were specified as descriptive decision rules.

### Candidate generation and receptor-matched controls within NK-RCPP

Target-sequence-conditioned peptide generation used PepMLM [10] through the PepMLM Google Colab notebook. The ChatterjeeLab/PepMLM-650M checkpoint and its tokenizer were loaded using Hugging Face AutoModelForMaskedLM and AutoTokenizer. NK1R, NK2R and NK3R sequences were used as separate target inputs. For each receptor, the requested peptide length was set to 10 residues (peptide_length = 10) and the number of generated candidates to 32 (num_binders = 32); the default top-k value was retained (top_k = 3). The notebook appended 10 mask tokens to the target sequence and obtained predictions for all masked positions in one model forward pass. At each position, the 3 highest-scoring token logits were converted to probabilities by softmax, and 1 token was sampled from the resulting categorical distribution. All peptide positions were sampled from the same fully masked input, rather than through iterative or autoregressive decoding. The logits were not temperature-scaled, equivalent to a sampling temperature of 1; the notebook exposed no separate temperature parameter. It selected CUDA when available and CPU otherwise, and did not set a fixed random seed.

The notebook calculated conditional pseudo-perplexity by concatenating the target and generated peptide sequences, masking each peptide position in turn and calculating the loss for the original token while retaining the other tokens. Pseudo-perplexity was the exponential of the mean peptide-position loss. Every generated candidate and its pseudo-perplexity were exported to CSV, without automatic ranking, filtering or deduplication. The public release provides pseudo-perplexity values for all 32 NK1R and 32 NK3R candidates; NK2R values and complete generation-run or subsequent selection records were unavailable. Pseudo-perplexity was not used in the joint prioritization rule.

The dataset contained 32 generated candidates per receptor. Most sequences had ten residues; NK2R-02 had eleven despite the requested length of ten, and its origin remained unresolved. Receptor-matched controls were Substance P (RPKPQQFFGLM) for NK1R, an NKA-derived peptide (HKTDSFVGLM) for NK2R and neurokinin B (DMHDFFVGLM) for NK3R. The original NK2R Colab export, executed notebook and checkpoint revisions, dependency versions and run-specific random-state information were unavailable.

### NK-RCPP on-target peptide-receptor docking and score extraction

Receptor inputs were NK1R from PDB 6E59 chain A (479 modelled residues), NK2R from a header-labelled AlphaFold monomer v2.0 model of UniProt P21452 (chain A, residues 1–398), and NK3R from a repository-described predicted model (chain B, 282 modelled residues within numbering 76–369; model source/version unrecorded). The NK1R input retains the glycogen-synthase fusion construct of the antagonist-bound 6E59 structure. Receptor preparation details and generation input sequences were unavailable in the supplied records.

The documented docking protocol used RAPiDock [13] and Rosetta REF15 [14], with ten poses per peptide, 16 diffusion steps and the local model. Repository settings specify rapidock_local.pt, conformation_partial = 1:1:1 and batch_size = 4. The on-target tables contain ten finite scores for each of the 99 study complexes, with rank-1 equal to the minimum in every case. We retained this minimum and summarized within-complex score dispersion by the range and sample standard deviation. The median within-complex range was 8,167.76 for NK1R, 1,158.96 for NK2R and 3,832.51 for NK3R, with 33 complexes including the control per receptor. The neighbour table provides rank1_score and best_score, which coincide for every stored complex, but lacks the ten underlying pose scores. Raw coordinates and preparation logs were unavailable for checking pose geometry, terminal modifications, protonation or disulfide assignments. We defined the on-target score difference as

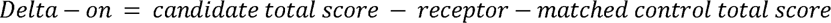

Delta-on < 0 denotes a lower on-target REF15 total model score than the receptor-matched control. Comparisons were confined to candidates and controls within the same receptor system. REF15 values are total model scores rather than binding free energies or calibrated affinities.

### NK-RCPP structural-neighbour panel construction

The repository workflow queried receptor structures against the Foldseek web service in combined 3Di and amino-acid mode [28]. The supplied search script listed 9 supported databases: afdb50, BFVD, afdb-swissprot, afdb-proteome, bfmd, mgnify_esm30, pdb100, cath50 and gmgcl_id. Its command-line default selected pdb100, afdb50, afdb-swissprot and afdb-proteome. Subsequent packaged processing used the PDB100 result table and wrote target sequences to FASTA. The processing script specifies CD-HIT [29] at sequence identity 0.50 when available, otherwise greedy 3-mer Jaccard clustering at 0.50. These are different similarity measures, not interchangeable identity thresholds; the available logs do not identify which branch was executed. Representative PDB entries were downloaded. The Foldseek-matched chain was extracted for docking, with the first chain used when the match could not be resolved.

Table S3 contains 66 NK1R-associated, 27 NK2R-associated and 60 NK3R-associated structures. Their identifiers match the released Foldseek hit tables and the complete panel used for each evaluable candidate. The panels include receptor-related structures, unrelated enzymes and the histamine H1 receptor–lysozyme chimera 3rze_A in the NK1R and NK3R sets. We used these heterogeneous structures as comparison panels. Raw search archives, the Foldseek server/database snapshot and final filtering logs were unavailable.

### Matched structural-neighbour scores and panel completeness

Candidate and receptor-matched control scores were paired on the same structural neighbours. We mapped sequences to canonical peptide codes before joining per-complex scores because NK2R batch numbering differs from the candidate-code system. This produced 4,518 paired rows over 82 candidates. Counts of common neighbours, counts with candidate-minus-control score > 0, and mean and median paired differences reproduced the candidate aggregates within floating-point rounding below 10 ¹². The source field frac_safer was termed the neighbour-score fraction and defined as

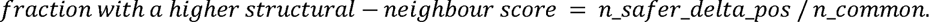

The neighbour-score fraction summarizes the direction of paired candidate-control score differences across structural contexts. The comparison-count proxy was n_common divided by the receptor-specific number of Table S3 rows. Identity-set checks confirmed that all 82 evaluable candidates covered their full 66-, 27- or 60-member panel, giving a proxy of 1.00. The ≥0.90 condition therefore served as a completeness check without further reducing this subset. Candidates without neighbour summaries remained unclassified under the joint rule.

### NK-RCPP prioritization rules and sensitivity analysis

The primary rule required Delta-on < 0 and a neighbour-score fraction > 0.50, with comparison-count proxy ≥0.90 as a completeness check. A higher-threshold sensitivity rule retained the completeness check and required a fraction ≥0.75. Rules were applied separately within each receptor. We also counted evaluable candidates at fraction cutoffs from >0.50 to >0.75 in increments of 0.05. For NK2R, the lower count bound was the observed evaluable-case count; the upper bound added every candidate with missing neighbour summaries and Delta-on < 0. Candidate ordering in figures prioritized the neighbour-score fraction and then Delta-on where relevant; complete on-target rankings were shown separately.

Joint and on-target-only selection were compared within the same 82 candidates with neighbour summaries. The threshold comparator retained all evaluable candidates with Delta-on < 0, and shortlist reduction was calculated relative to this pool. For the equal-size comparison, we ranked evaluable candidates by ascending Delta-on within each receptor and selected the top three NK1R, ten NK2R and three NK3R candidates, matching the joint-rule quotas. Candidate identifier would resolve ties. We counted the intersection of the two sets within each receptor and summed the intersections across receptors. Candidates missing neighbour summaries were excluded from both complete-case comparisons. Activity labels were unavailable for the 96 candidates, and the comparison endpoint was candidate selection. Missing NK2R pseudo-perplexity values precluded a complete receptor-wide comparison with the native generator ranking; the available NK1R/NK3R values were not used to redefine the selection rules.

### Sequence-frequency summaries

Generated sequences of exactly ten residues were aligned by position without gaps. For each receptor and position, amino-acid frequency was the residue count divided by the number of included sequences. Heatmaps used the fixed order ACDEFGHIKLMNPQRSTVWY. **Fig. 4** uses a shared scale of 0–0.60, while **Supplementary Fig. S9** uses 0–0.50, clipping larger frequencies at that colour limit without changing the tabulated values. A consensus character was shown when the most frequent residue at a position reached at least 0.30. The eleven-residue NK2R-02 was excluded.

### Exploratory NK2R comparison and sequence-space projection

The exploratory NK2R comparison comprised ten competition peptides with workbook-reported EC50 values and five random controls at each of six, nine, ten and fifteen residues. The source workbook was available locally; its formal public source and originating assay protocols/raw records were unavailable. The standardized score was the peptide score minus the same-length random-control mean, divided by that group’s sample standard deviation. NK2R-02 remained in raw-score analyses but was excluded from standardization because no eleven-residue baseline was available. The ten-residue random-control pool was constructed from NK2R-01 and the NKA-derived control. Group comparisons used two-sided asymptotic Mann–Whitney tests with pooled average ranks, tie-corrected variance and a 0.5 continuity correction: 10 versus 32 for raw scores, 10 versus 18 for neighbour-score fractions and 10 versus 31 for standardized scores. Competition peptides were the first group for U. Signed rank-biserial correlation was twice the proportion of cross-group pairs with higher competition values (ties counted half), minus one; positive values indicate higher competition-group scores. Spearman rho used average ranks and a two-sided Student-t approximation with n−2 degrees of freedom. All tests were exploratory and uncorrected for multiplicity.

The sequence comparison set combined the first 1,000 entries of the supplied 2025 peptide-design competition workbook with 96 study candidates and three receptor-matched controls. Standardization and deduplication, preferentially retaining study entries, yielded 1,048 unique sequences: 949 leaderboard and 99 study entries. Five study sequences also occurred among the original leaderboard entries. The embedding script specifies ESM-2 esm2_t33_650M_UR50D [25], layer 33, with mean pooling over residue tokens excluding beginning/end tokens. The projection script specifies L2 normalization followed by UMAP [26] with n_neighbors = 30, min_dist = 0.30, metric = cosine, n_components = 2 and random_state = 42. We reused the saved coordinates and calculated nearest-reference Euclidean distances against the 949 retained leaderboard entries in the two-dimensional projection, keeping the ten-candidate and 99-entry medians separate. The projection and distances were used for sequence visualization.

### Reproducibility and figure generation

Reanalysis used Python 3.12.14, pandas 3.0.1 and NumPy 2.3.5. We checked labels, identifiers, sequence lengths and panel sizes, reconstructed paired differences and candidate summaries, and compared them with the source tables. Statistical calculations used local standard-library implementations of the methods above, cross-checked by rank-sum/pair-count calculations and Student-t integration. Source-file SHA-256 values, validation tables, scripts and figure sources are included in the accompanying support archive. **Fig. 5** and **Supplementary Figs. S6–S8** were exported from editable SVG to 600-dpi PNG and LZW-compressed TIFF. The workflow raster and other figures were retained from the source archive. The seed 20260801 controls plotting jitter; the UMAP script specifies random_state = 42. Historical generation, docking, search and embedding dependency versions were unavailable.

## Data Availability

Candidate and receptor-matched control sequences, on-target scores, Foldseek structural-neighbour records, candidate aggregates, exploratory NK2R comparison data, length-control data and sequence-projection coordinates are available at https://github.com/HUST-NingKang-Lab/NK-RCPP. Figure sources and supplementary tables are included. The release at commit 4a04cab13476e30ba5cd1a24ea876ca6d9305998 contains per-complex neighbour scores and pair-level comparison tables. Table S3 is provided as figures/tables/TableS3_offtarget_proteins.csv. Validation tables and the figure sources associated with the reanalysis are provided in the accompanying support archive. Raw RAPiDock pose directories, the original NK2R Colab export and the source competition workbook are not deposited. Data and figures are released under CC BY 4.0, with third-party inputs subject to their original terms.

## Code Availability

Analysis and plotting code, workflow documentation and usage guidance are available at https://github.com/HUST-NingKang-Lab/NK-RCPP under the MIT licence. The scripts cover score extraction, paired structural-neighbour comparisons, candidate prioritization, exploratory NK2R comparisons and figure generation. The accompanying support archive contains the reanalysis and plotting sources used for this manuscript; repository README files provide usage instructions and file mappings for the original workflow. PepMLM, RAPiDock, Rosetta, Foldseek, CD-HIT, ESM-2 and UMAP are third-party resources described in Methods and remain subject to their respective licences and access conditions.

## Supplementary figures and legends

**Supplementary Fig. S1.**
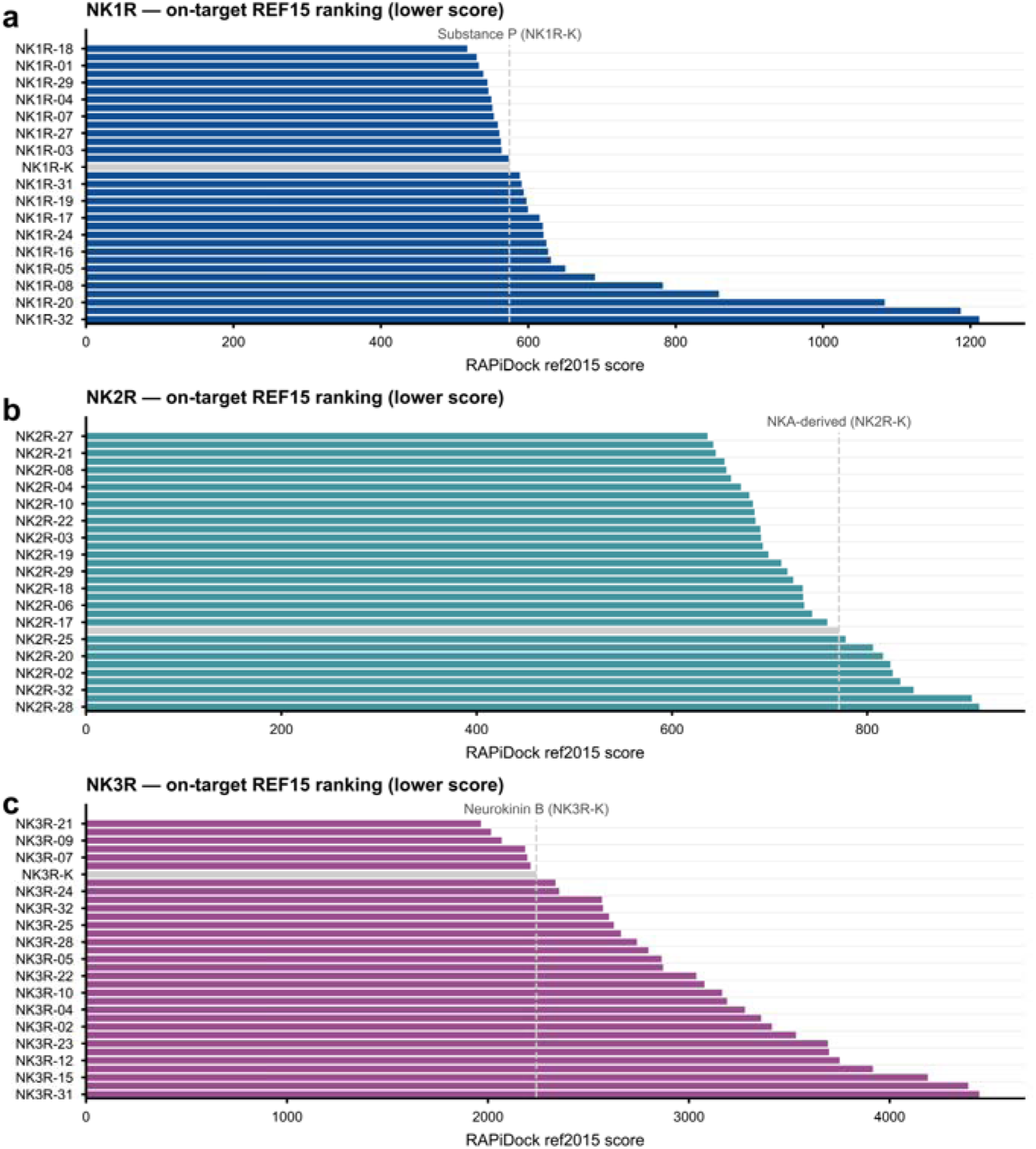
Complete on-target REF15 rankings. Complete within-receptor rankings by minimum REF15 total model score. a, NK1R candidates and the Substance P control. b, NK2R candidates and the NKA-derived control. c, NK3R candidates and the neurokinin B control. Lower scores appear toward the top. Each study complex has ten on-target scores, with rank-1 equal to the minimum in all 99 complexes. Rankings compare candidates with their receptor-matched control.

**Supplementary Fig. S2.**
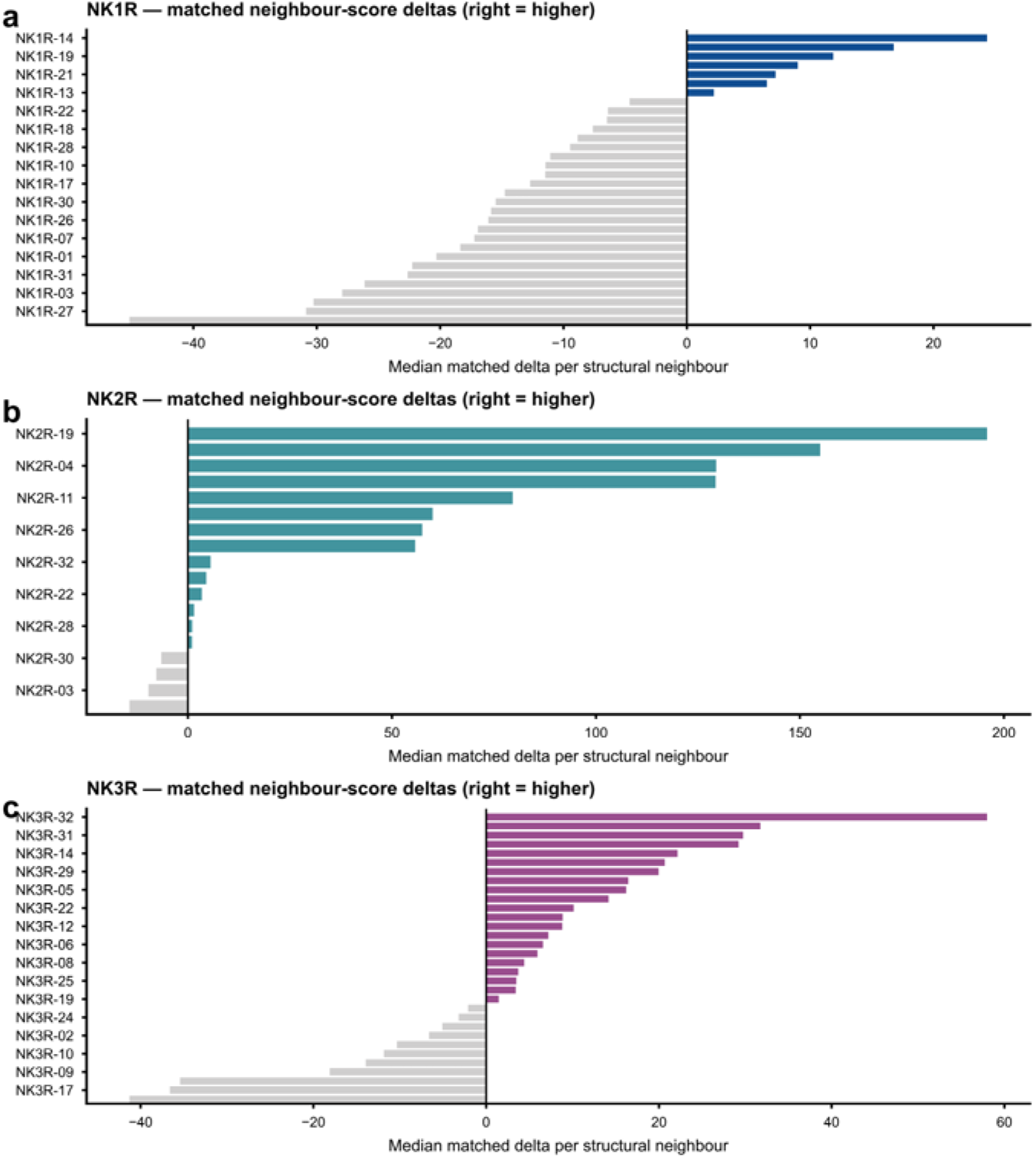
Matched structural-neighbour score differences. Median paired candidate-control score differences across structural-neighbour comparisons for a, NK1R; b, the evaluable NK2R candidates; and c, NK3R. Positive values indicate a higher candidate REF15 total model score than the matched control on the same structural neighbour.

**Supplementary Fig. S3.**
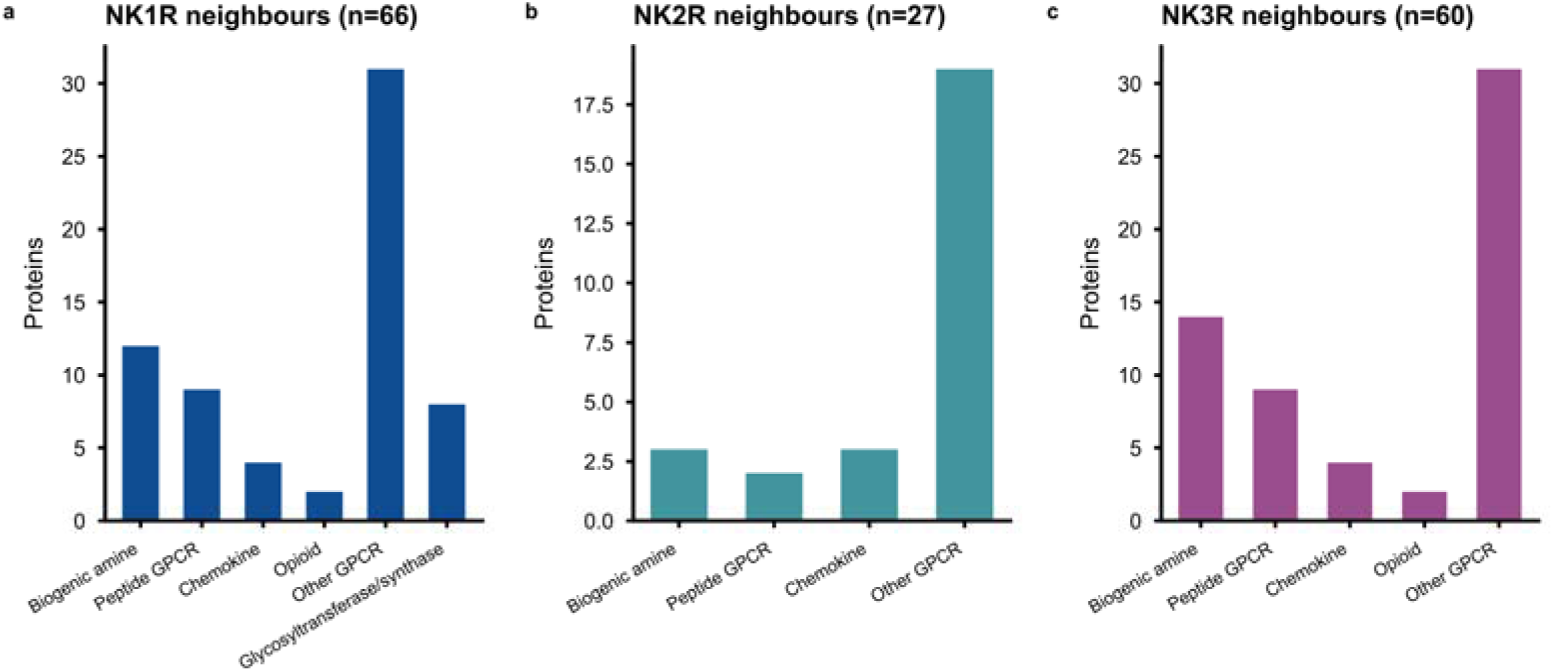
Structural-neighbour panel composition. Annotation composition of structural-neighbour panels for a, NK1R (n = 66); b, NK2R (n = 27); and c, NK3R (n = 60). Categories summarize the annotations of the comparison structures.

**Supplementary Fig. S4.**
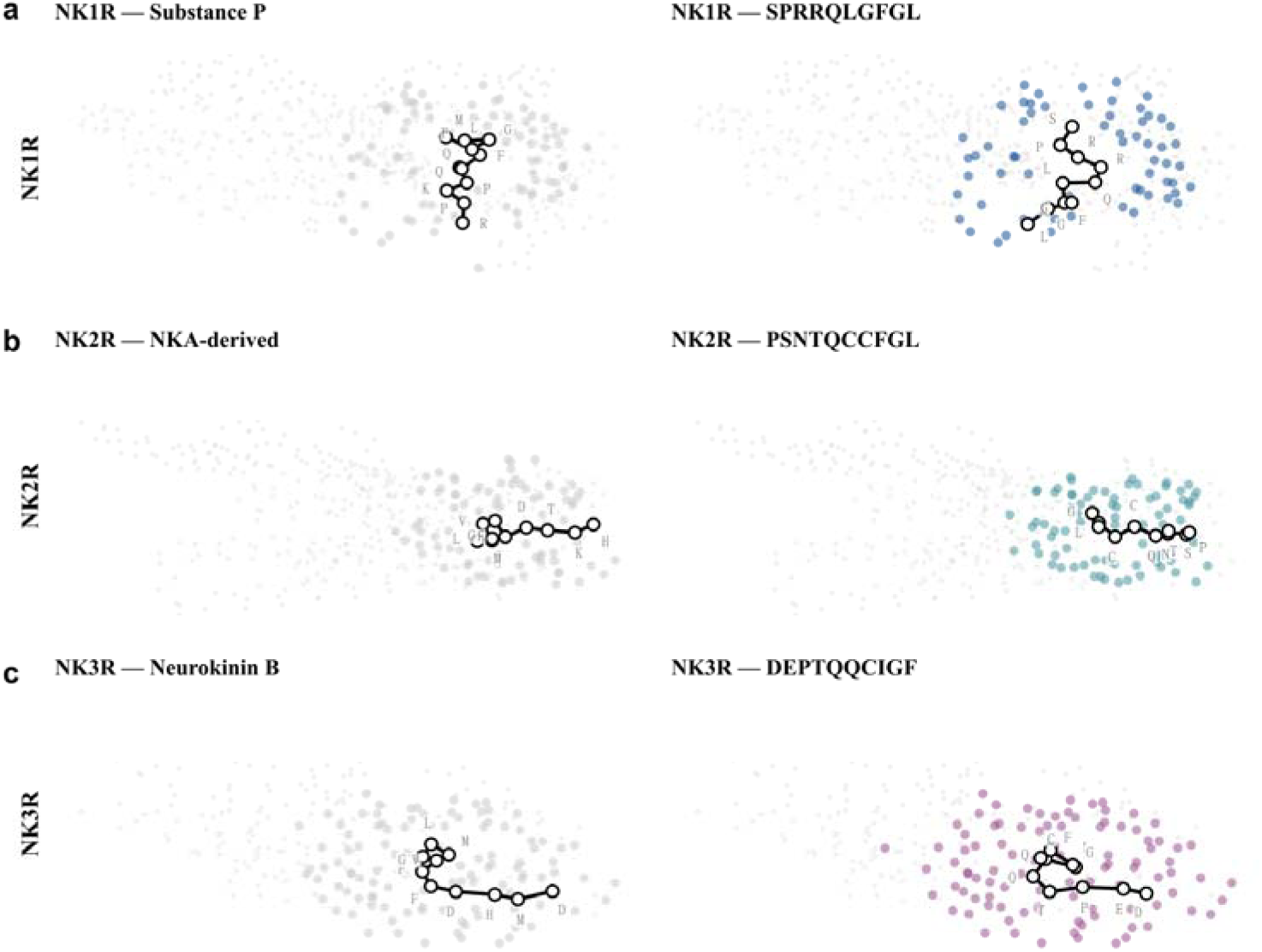
Representative archived peptide-pocket views. Representative archived peptide-pocket views. Receptor-matched control and candidate renderings for a, NK1R; b, NK2R; and c, NK3R. Panels reproduce the computational views retained in the source archive.

**Supplementary Fig. S5.**
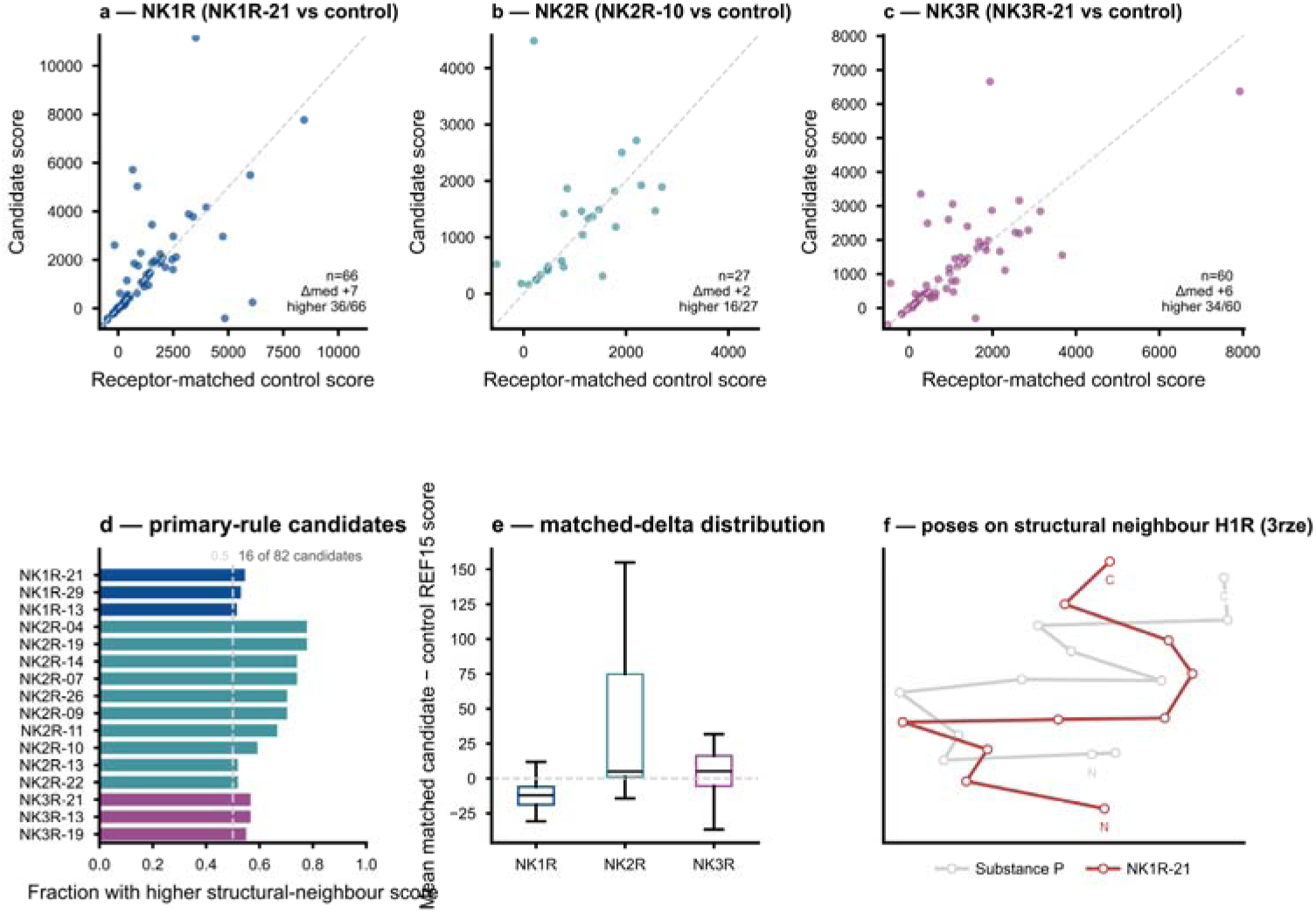
Candidate-control structural-neighbour comparisons. Paired candidate-control structural-neighbour scores. a–c, Candidate and matched control REF15 scores on common structural neighbours for one representative candidate per receptor; points above the diagonal have a higher candidate score. d, Neighbour-score fractions for primary-rule candidates. e, Distributions of paired score differences. f, Archived illustration of NK1R peptide traces on one structural neighbour.

**Supplementary Fig. S6.**
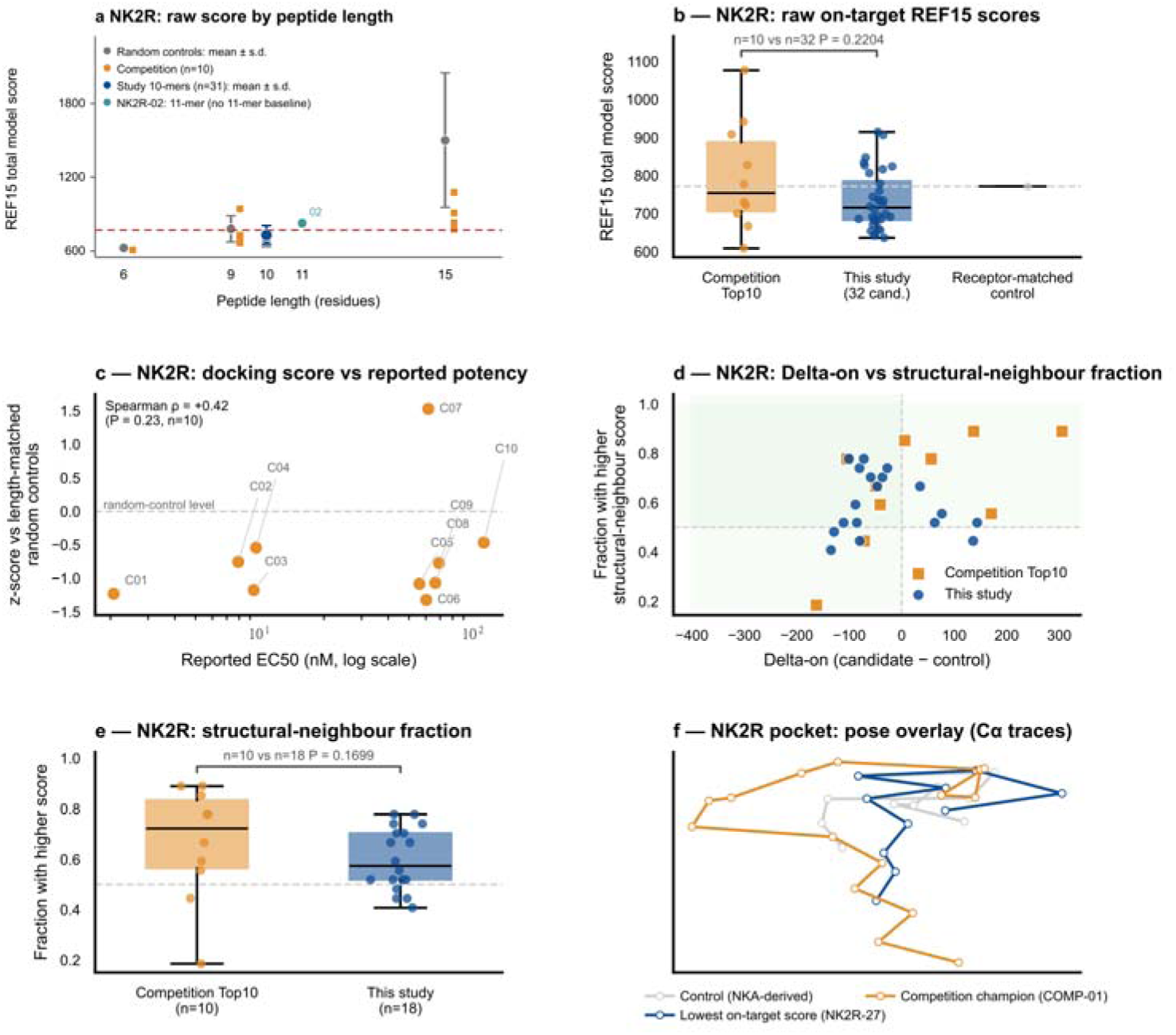
External NK2R competition comparison. Exploratory NK2R peptide comparison. a, Raw scores by peptide length: random controls (five per length, mean ± s.d.), competition peptides (n = 10), study ten-mers (n = 31, mean ± s.d.) and the eleven-residue NK2R-02, which lacks a matching baseline. The dashed line marks the receptor-matched control score. b, Raw-score distributions for all 32 study candidates and ten competition peptides (competition-first U = 202, rank-biserial r = +0.263, P = 0.2204). c, Standardized score versus workbook-reported EC50 (Spearman rho = +0.42, P = 0.23, n = 10). d, Delta-on versus neighbour-score fraction. e, Fractions for 18 evaluable study candidates and ten competition peptides (U = 119, r = +0.322, tie-corrected P = 0.1699). f, Archived NK2R trace overlay including NK2R-27, the candidate with the lowest on-target score. Tests are exploratory, two-sided and uncorrected for multiplicity. Random-control residue pools differ between lengths.

**Supplementary Fig. S7.**
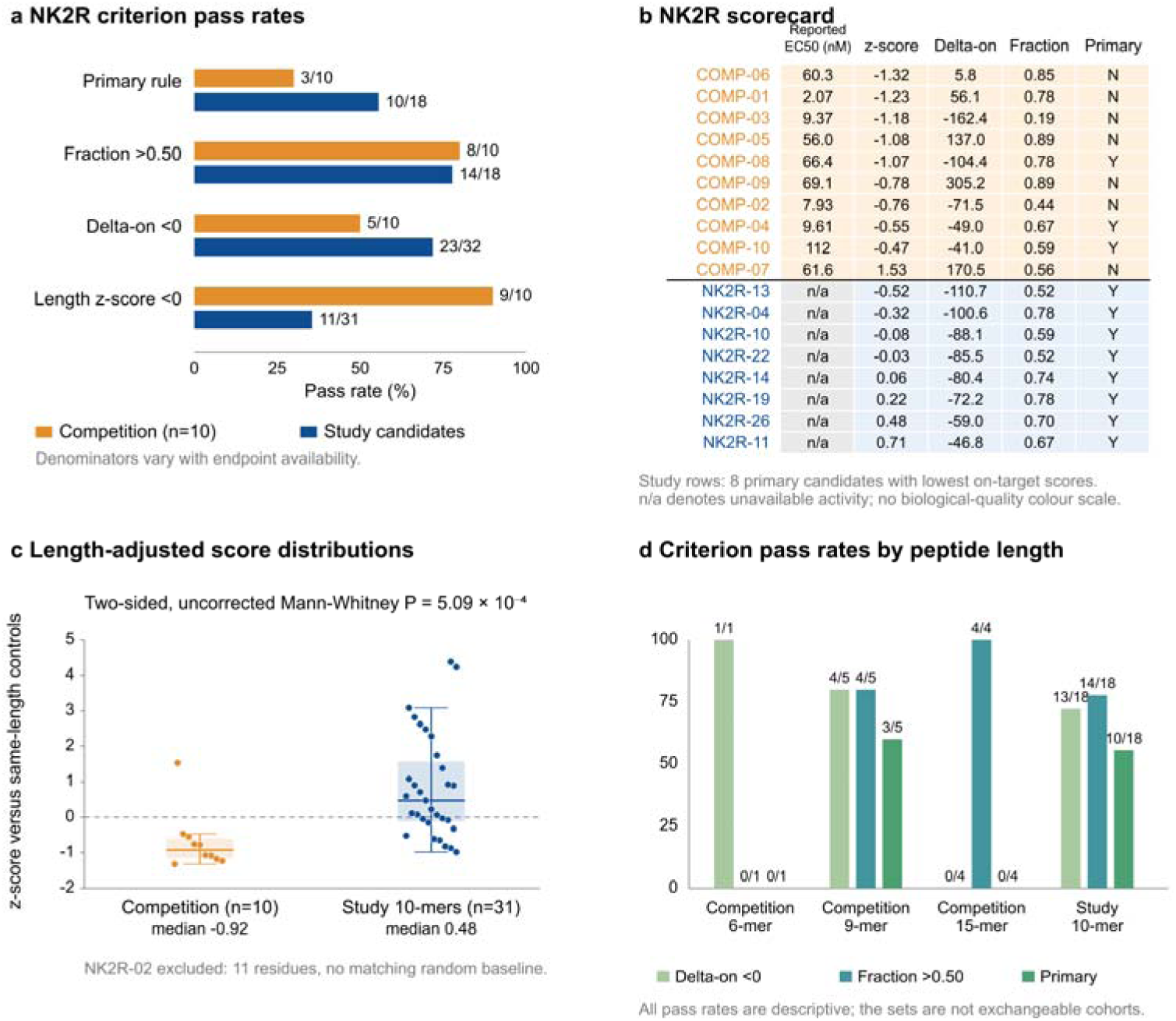
Descriptive NK2R benchmark summaries. Descriptive NK2R comparison summaries. a, Pass counts with endpoint-specific denominators: competition n = 10; study n = 32 for Delta-on, n = 18 for neighbour-score fractions and the joint rule, and n = 31 for standardized scores. b, Scorecard for ten competition peptides and the eight primary-rule study candidates with the lowest on-target scores; n/a denotes unavailable activity. c, Standardized scores for ten competition peptides and 31 study ten-mers, excluding NK2R-02 because no eleven-residue baseline was available (competition-first U = 40, rank-biserial r = −0.742, P = 5.09 × 10). Boxes show medians and quartiles; whiskers extend to the furthest observations within 1.5 interquartile ranges, and all observations are shown. d, Pass counts by length. Primary denotes Delta-on < 0 and fraction > 0.50. Tests are exploratory, two-sided and uncorrected for multiplicity.

**Supplementary Fig. S8.**
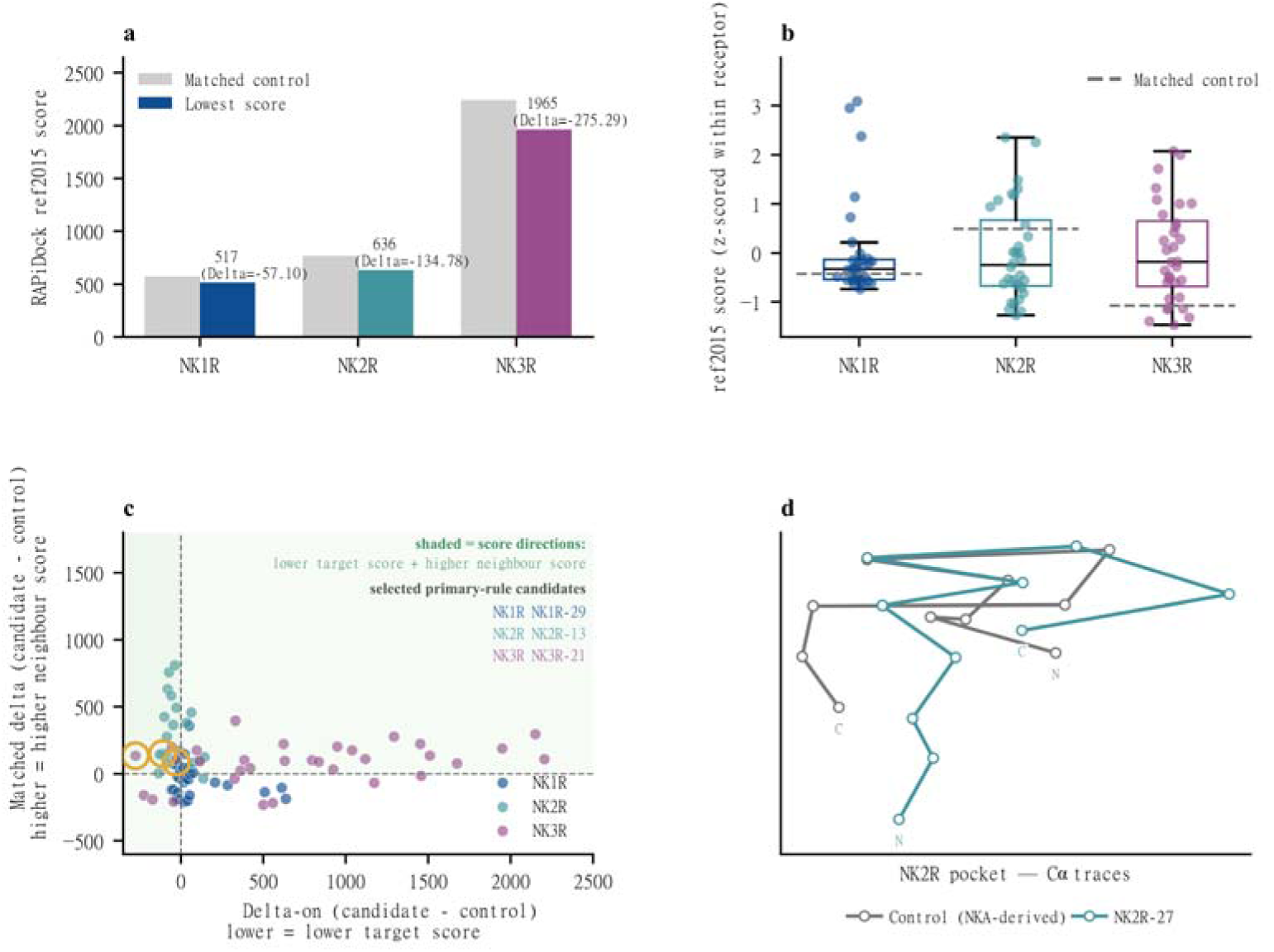
Extended docking-score diagnostics. Extended docking-score summaries. a–b, Receptor-matched control and lowest candidate scores, candidate-minus-control differences, and within-receptor standardized score distributions. Standardization uses the mean and sample standard deviation of the 32 candidates within each receptor. c–d, Joint Delta-on and paired structural-neighbour score-difference landscape, with an archived NK2R peptide trace overlay. Shading marks the direction of the computational selection criteria.

**Supplementary Fig. S9.**
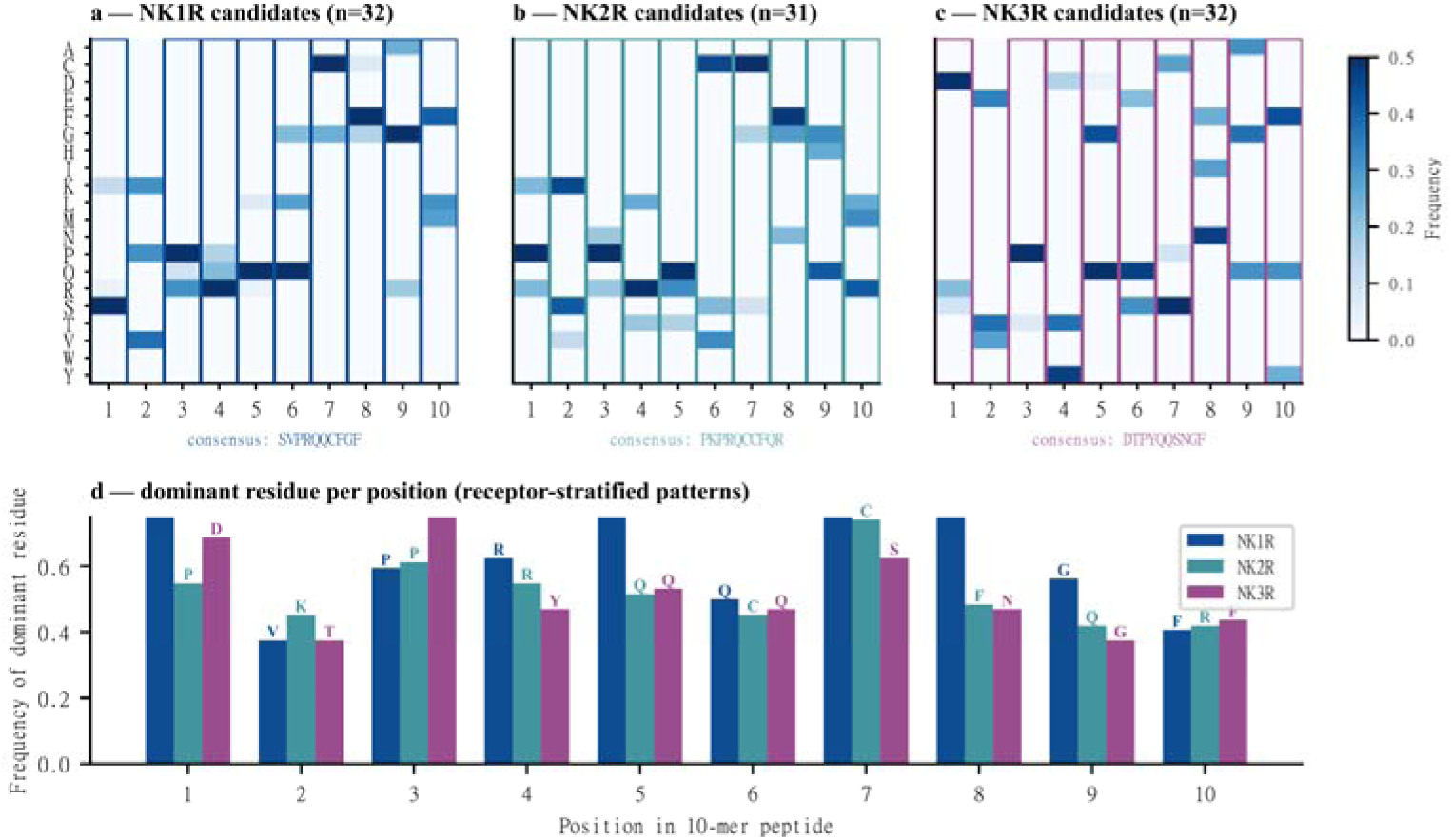
Extended sequence-frequency summaries. Extended sequence-frequency summaries. a–c, Position-wise amino-acid frequencies for exact ten-residue NK1R, NK2R and NK3R candidates; the eleven-residue NK2R-02 was excluded. d, Most frequent residue at each aligned position, stratified by receptor.

## References

1. Sriram, K. and P.A. Insel, G Protein-Coupled Receptors as Targets for Approved Drugs: How Many Targets and How Many Drugs? Molecular Pharmacology, 2018. 93(4): p. 251–258.

2. Steinhoff, M.S., et al., Tachykinins and Their Receptors: Contributions to Physiological Control and the Mechanisms of Disease. Physiological Reviews, 2014. 94(1): p. 265–301.

3. Hesketh, P.J., et al., The Oral Neurokinin-1 Antagonist Aprepitant for the Prevention of Chemotherapy-Induced Nausea and Vomiting: A Multinational, Randomized, Double-Blind, Placebo-Controlled Trial in Patients Receiving High-Dose Cisplatin—The Aprepitant Protocol 052 Study Group. Journal of Clinical Oncology, 2003. 21(22): p. 4112–4119.

4. Lederman, S., et al., Fezolinetant for treatment of moderate-to-severe vasomotor symptoms associated with menopause (SKYLIGHT 1): a phase 3 randomised controlled study. The Lancet, 2023. 401(10382): p. 1091–1102.

5. Thom, C., et al., Structures of neurokinin 1 receptor in complex with Gq and Gs proteins reveal substance P binding mode and unique activation features. Science Advances, 2021. 7(50): p. eabk2872.

6. Sun, W., et al., Structural insights into neurokinin 3 receptor activation by endogenous and analogue peptide agonists. Cell Discovery, 2023. 9(1): p. 66.

7. Petersen, J.E., et al., Molecular determinants of neuropeptide-mediated activation mechanisms in tachykinin NK1 and NK2 receptors. Journal of Biological Chemistry, 2024. 300(12): p. 107948.

8. Rupniak, N.M.J., et al., Affinity, potency, efficacy, and selectivity of neurokinin A analogs at human recombinant NK2 and NK1 receptors. PLOS ONE, 2018. 13(10): p. e0205894.

9. Muttenthaler, M., et al., Trends in peptide drug discovery. Nature Reviews Drug Discovery, 2021. 20(4): p. 309–325.

10. Chen, L.T., et al., Target sequence-conditioned design of peptide binders using masked language modeling. Nature Biotechnology, 2026. 44(6): p. 1002–1010.

11. Madani, A., et al., Large language models generate functional protein sequences across diverse families. Nature Biotechnology, 2023. 41(8): p. 1099–1106.

12. Ciemny, M., et al., Protein–peptide docking: opportunities and challenges. Drug Discovery Today, 2018. 23(8): p. 1530–1537.

13. Zhao, H., et al., Protein–peptide docking with a rational and accurate diffusion generative model. Nature Machine Intelligence, 2025. 7(8): p. 1308–1321.

14. Alford, R.F., et al., The Rosetta All-Atom Energy Function for Macromolecular Modeling and Design. Journal of Chemical Theory and Computation, 2017. 13(6): p. 3031–3048.

15. Tsaban, T., et al., Harnessing protein folding neural networks for peptide–protein docking. Nature Communications, 2022. 13(1): p. 176.

16. Junker, H. and C.T. Schoeder, Assessment of generative de novo peptide design methods for G protein-coupled receptors. PLOS One, 2026. 21(8): p. e0355549.

17. Repecka, D., et al., Expanding functional protein sequence spaces using generative adversarial networks. Nature Machine Intelligence, 2021. 3(4): p. 324–333.

18. Dauparas, J., et al., Robust deep learning–based protein sequence design using ProteinMPNN. Science, 2022. 378(6615): p. 49–56.

19. Ingraham, J.B., et al., Illuminating protein space with a programmable generative model. Nature, 2023. 623(7989): p. 1070–1078.

20. Cao, L., et al., Design of protein-binding proteins from the target structure alone. Nature, 2022. 605(7910): p. 551–560.

21. Watson, J.L., et al., De novo design of protein structure and function with RFdiffusion. Nature, 2023. 620(7976): p. 1089–1100.

22. Vázquez Torres, S., et al., De novo design of high-affinity binders of bioactive helical peptides. Nature, 2024. 626(7998): p. 435–442.

23. Pacesa, M., et al., One-shot design of functional protein binders with BindCraft. Nature, 2025. 646(8084): p. 483–492.

24. Bhat, S., et al., De novo design of peptide binders to conformationally diverse targets with contrastive language modeling. Science Advances, 2025. 11(4): p. eadr8638.

25. Lin, Z., et al., Evolutionary-scale prediction of atomic-level protein structure with a language model. Science, 2023. 379(6637): p. 1123–1130.

26. McInnes, L., et al., UMAP: Uniform Manifold Approximation and Projection. Journal of Open Source Software, 2018. 3(29): p. 861.

27. Kenakin, T. and A. Christopoulos, Signalling bias in new drug discovery: detection, quantification and therapeutic impact. Nature Reviews Drug Discovery, 2013. 12(3): p. 205–216.

28. van Kempen, M., et al., Fast and accurate protein structure search with Foldseek. Nature Biotechnology, 2024. 42(2): p. 243–246.

29. Li, W. and A. Godzik, Cd-hit: a fast program for clustering and comparing large sets of protein or nucleotide sequences. Bioinformatics, 2006. 22(13): p. 1658–1659.

