## supplementary figures for "NK-RCPP: a receptor-conditioned computational workflow for prioritizing neurokinin-receptor peptide candidates"

**NK-RCPP Supplementary Figures**

### Supplementary Fig. S1 | Complete on-target REF15 rankings

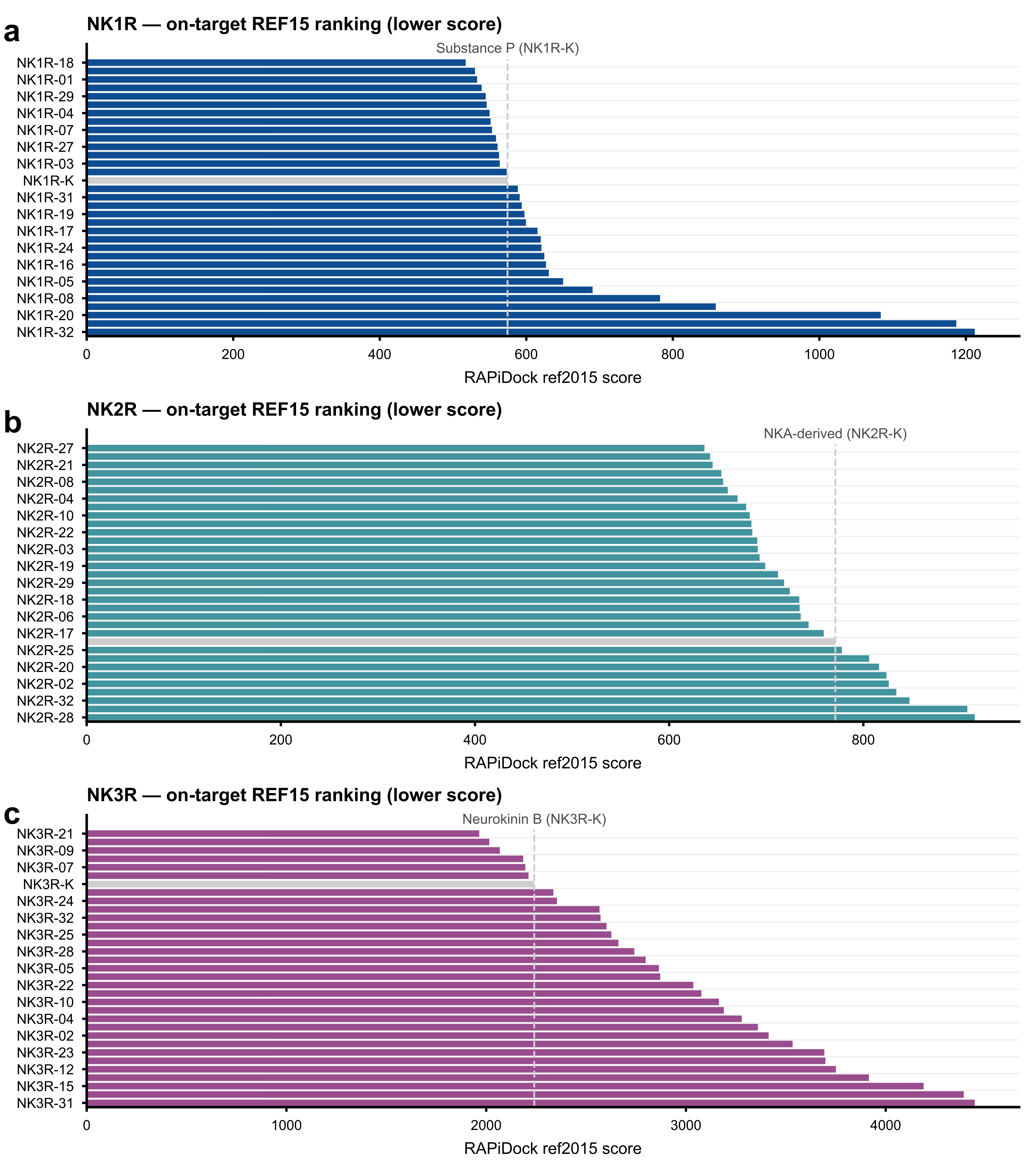

**Supplementary Fig. S1** | Complete within-receptor rankings by minimum REF15 total model score. a, NK1R candidates and the Substance P control. b, NK2R candidates and the NKA-derived control. c, NK3R candidates and the neurokinin B control. Lower scores appear toward the top. Each study complex has ten on-target scores, with rank-1 equal to the minimum in all 99 complexes. Rankings compare candidates with their receptor-matched control.

### Supplementary Fig. S2 | Matched structural-neighbour score differences

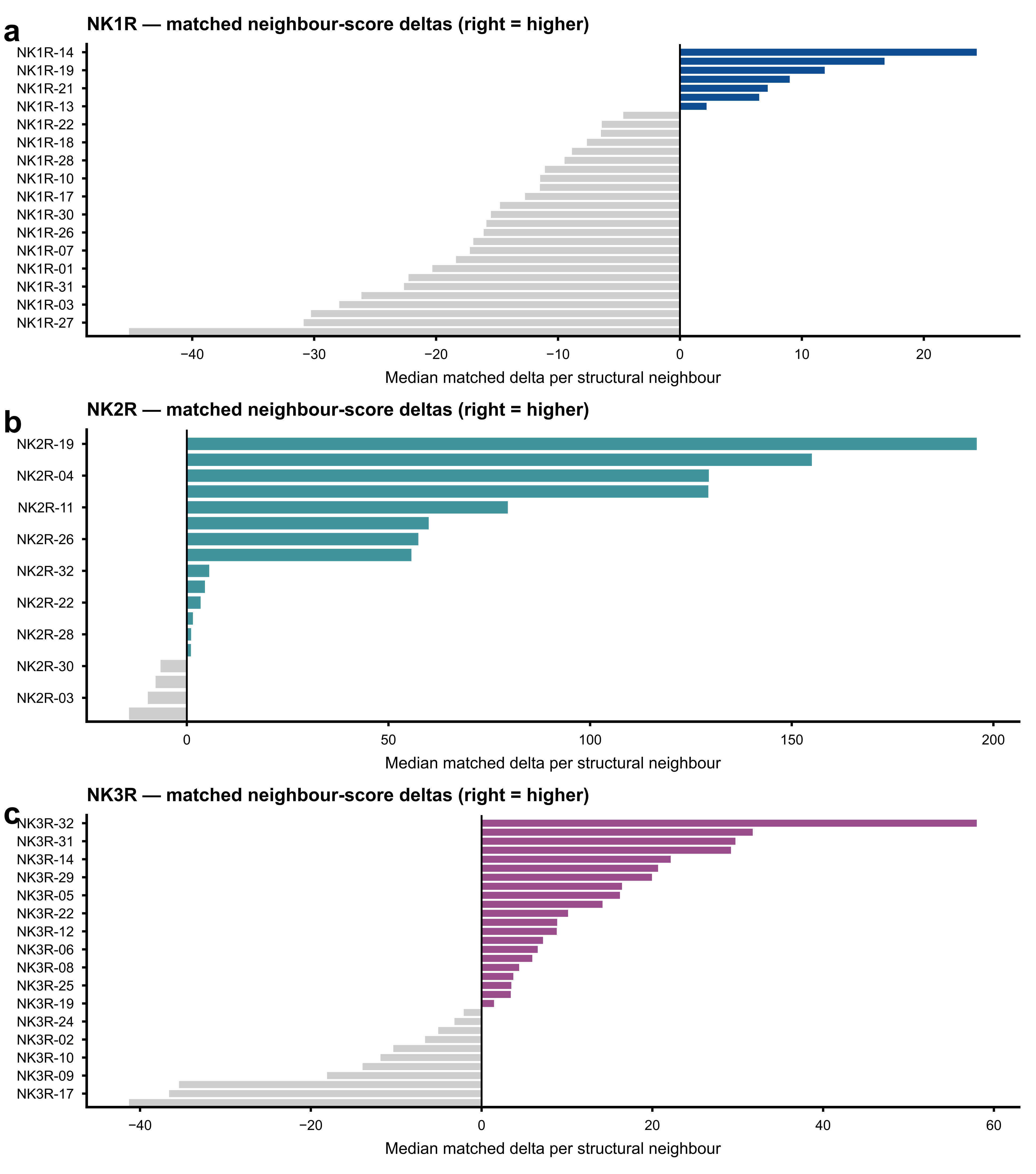

Supplementary Fig. S2 | Median paired candidate-control score differences across structural-neighbour comparisons for a, NK1R; b, the evaluable NK2R candidates; and c, NK3R. Positive values indicate a higher candidate REF15 total model score than the matched control on the same structural neighbour.

### Supplementary Fig. S3 | Structural-neighbour panel composition

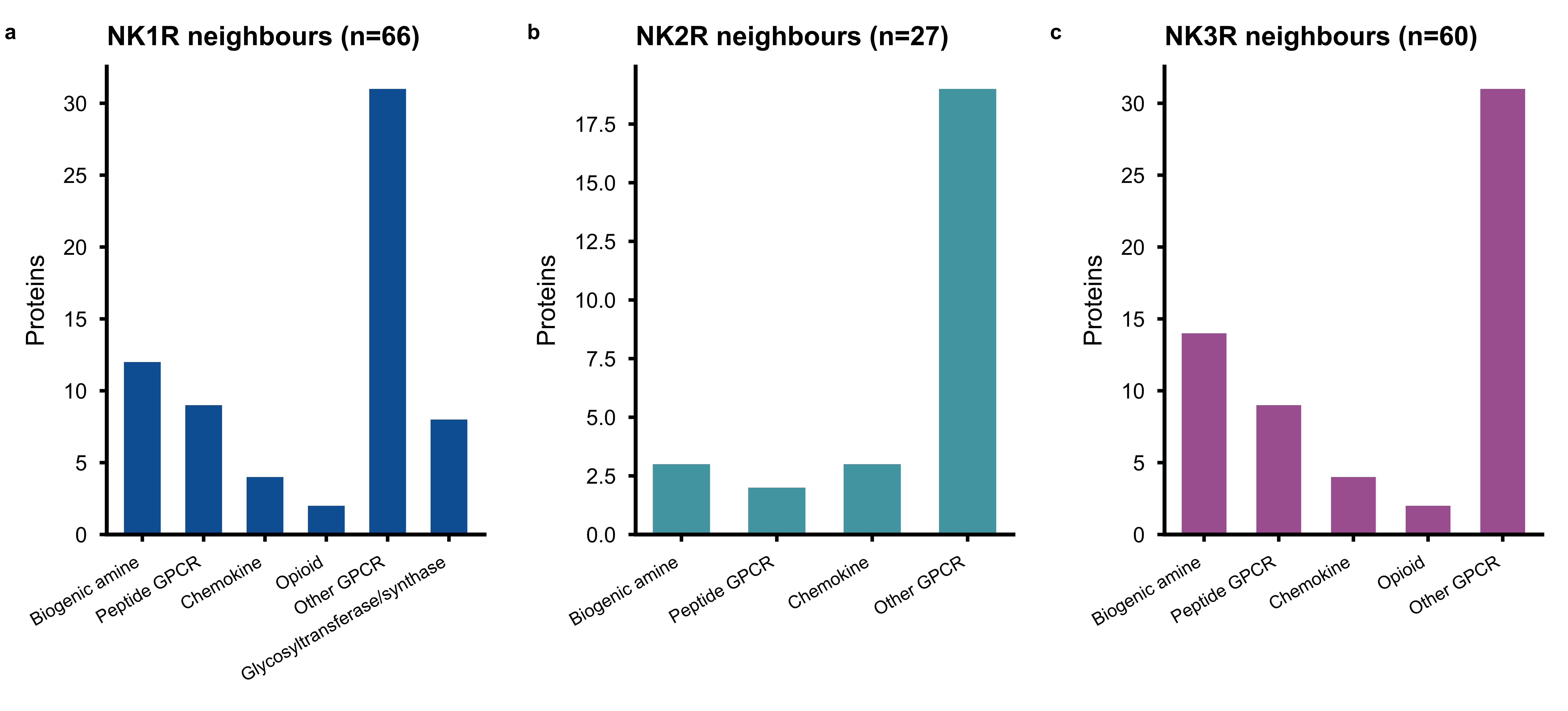

Supplementary Fig. S3 | Annotation composition of structural-neighbour panels for a, NK1R (n = 66); b, NK2R (n = 27); and c, NK3R (n = 60). Categories summarize the annotations of the comparison structures.

### Supplementary Fig. S4 | Representative archived peptide-pocket views

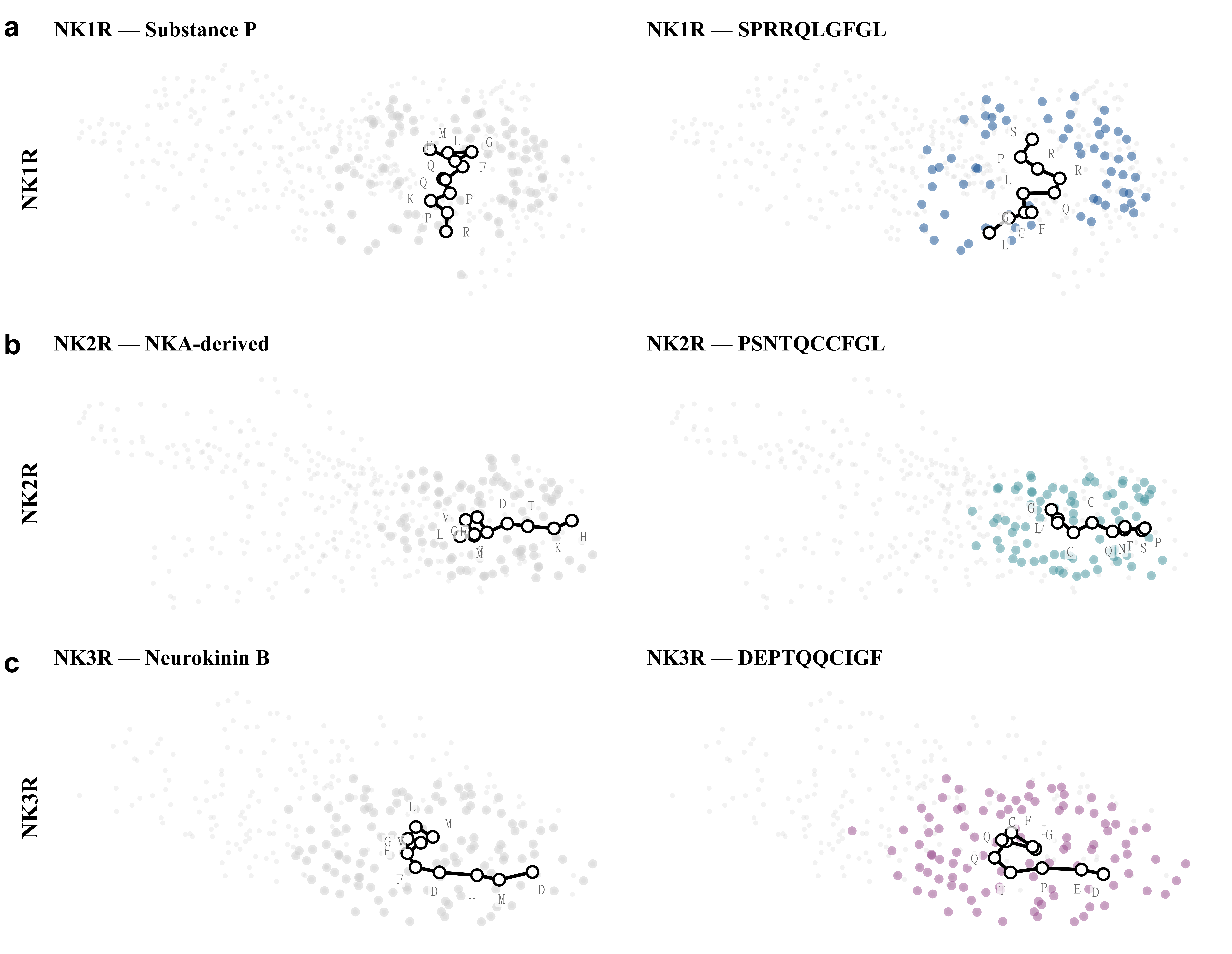

Supplementary Fig. S4 | Representative archived peptide-pocket views. Receptor-matched control and candidate renderings for a, NK1R; b, NK2R; and c, NK3R. Panels reproduce the computational views retained in the source archive.

### Supplementary Fig. S5 | Candidate-control structural-neighbour comparisons

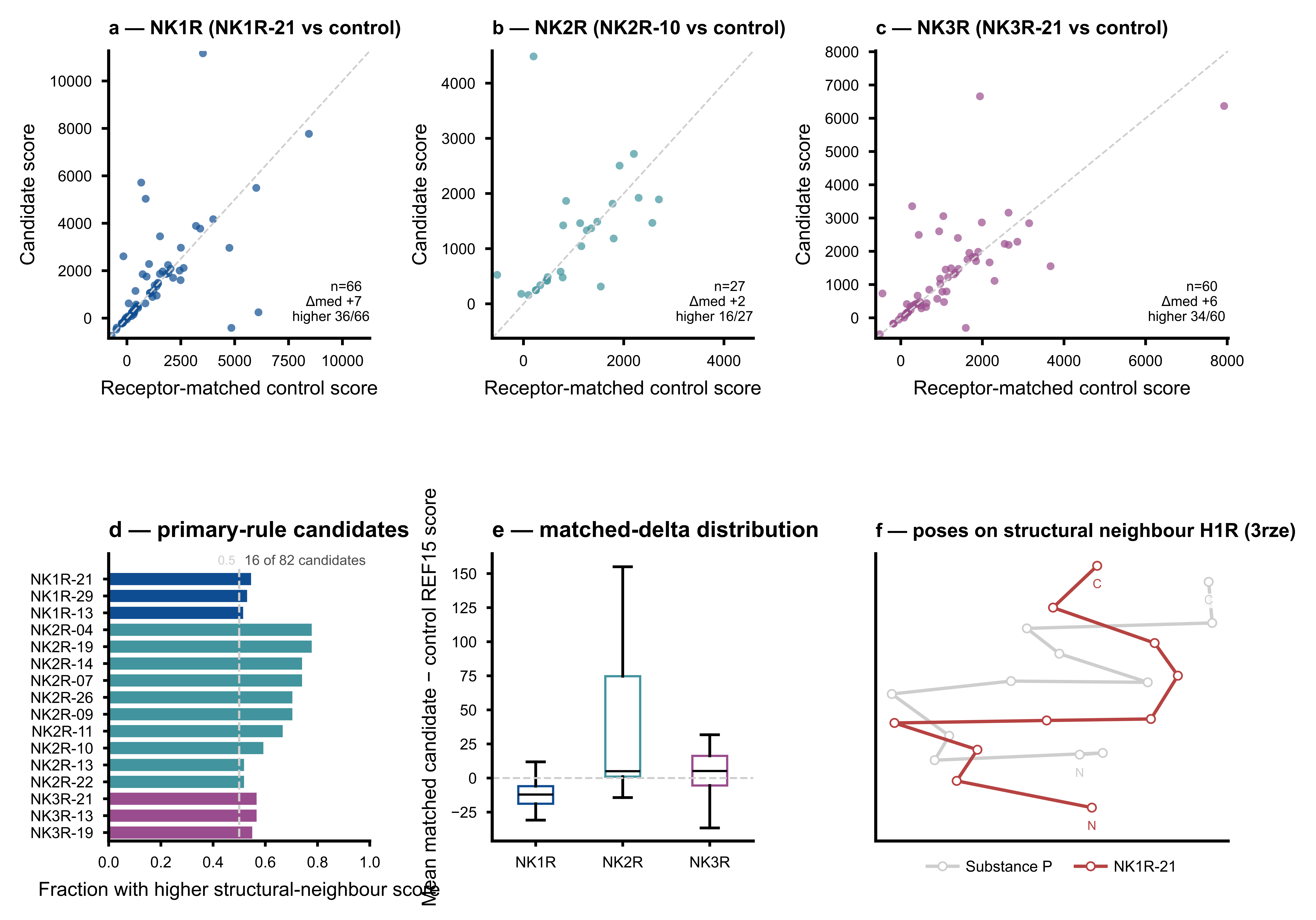

**Supplementary Fig. S5** | Paired candidate-control structural-neighbour scores. a–c, Candidate and matched control REF15 scores on common structural neighbours for one representative candidate per receptor; points above the diagonal have a higher candidate score. d, Neighbour-score fractions for primary-rule candidates. e, Distributions of paired score differences. f, Archived illustration of NK1R peptide traces on one structural neighbour.

### Supplementary Fig. S6 | External NK2R competition comparison

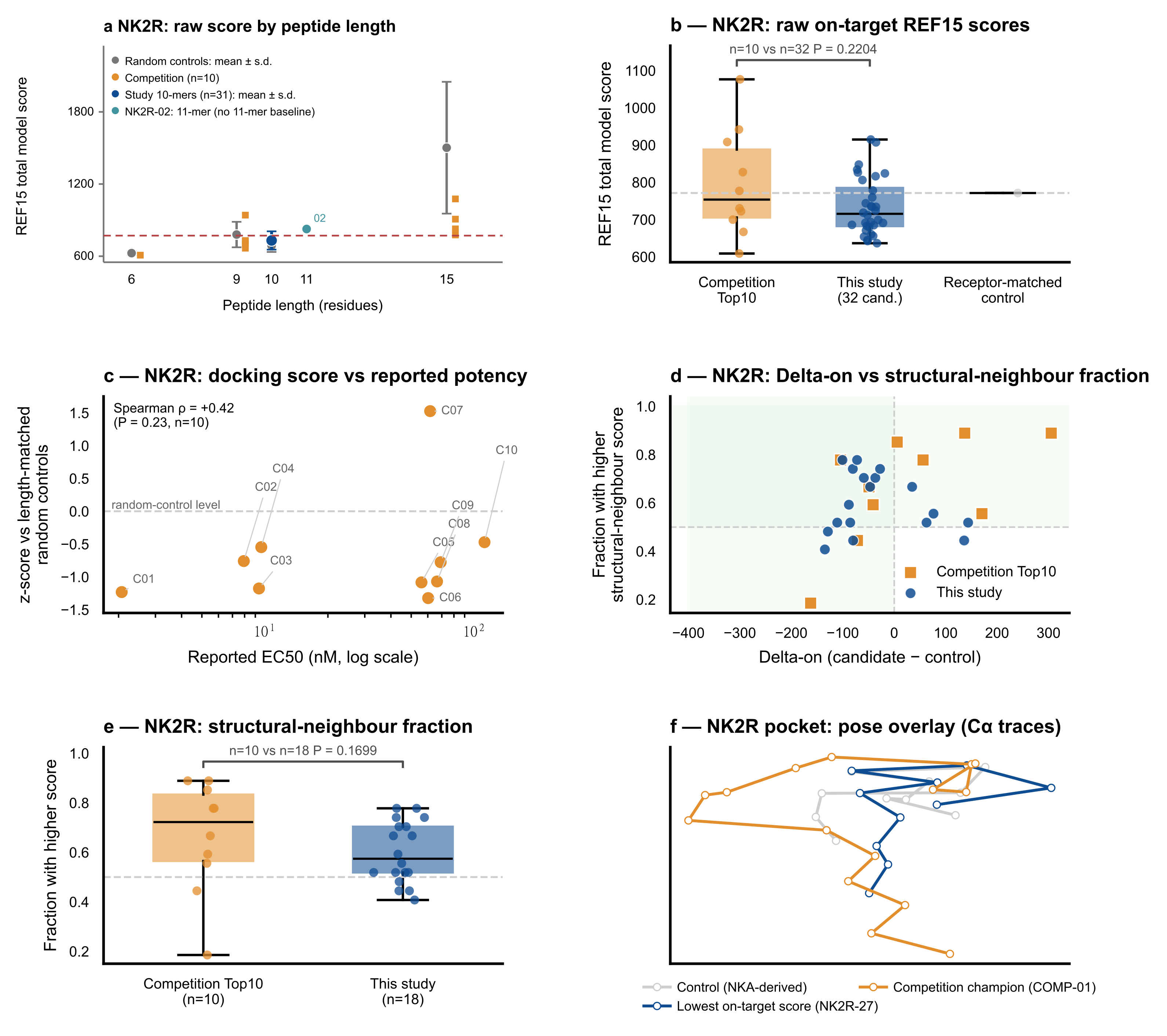

**Supplementary Fig. S6** | Exploratory NK2R peptide comparison. a, Raw scores by peptide length: random controls (five per length, mean ± s.d.), competition peptides (n = 10), study ten-mers (n = 31, mean ± s.d.) and the eleven-residue NK2R-02, which lacks a matching baseline. The dashed line marks the receptor-matched control score. b, Raw-score distributions for all 32 study candidates and ten competition peptides (competition-first U = 202, rank-biserial r = +0.263, P = 0.2204). c, Standardized score versus workbook-reported EC50 (Spearman rho = +0.42, P = 0.23, n = 10). d, Delta-on versus neighbour-score fraction. e, Fractions for 18 evaluable study candidates and ten competition peptides (U = 119, r = +0.322, tie-corrected P = 0.1699). f, Archived NK2R trace overlay including NK2R-27, the candidate with the lowest on-target score. Tests are exploratory, two-sided and uncorrected for multiplicity. Random-control residue pools differ between lengths.

### Supplementary Fig. S7 | Descriptive NK2R benchmark summaries

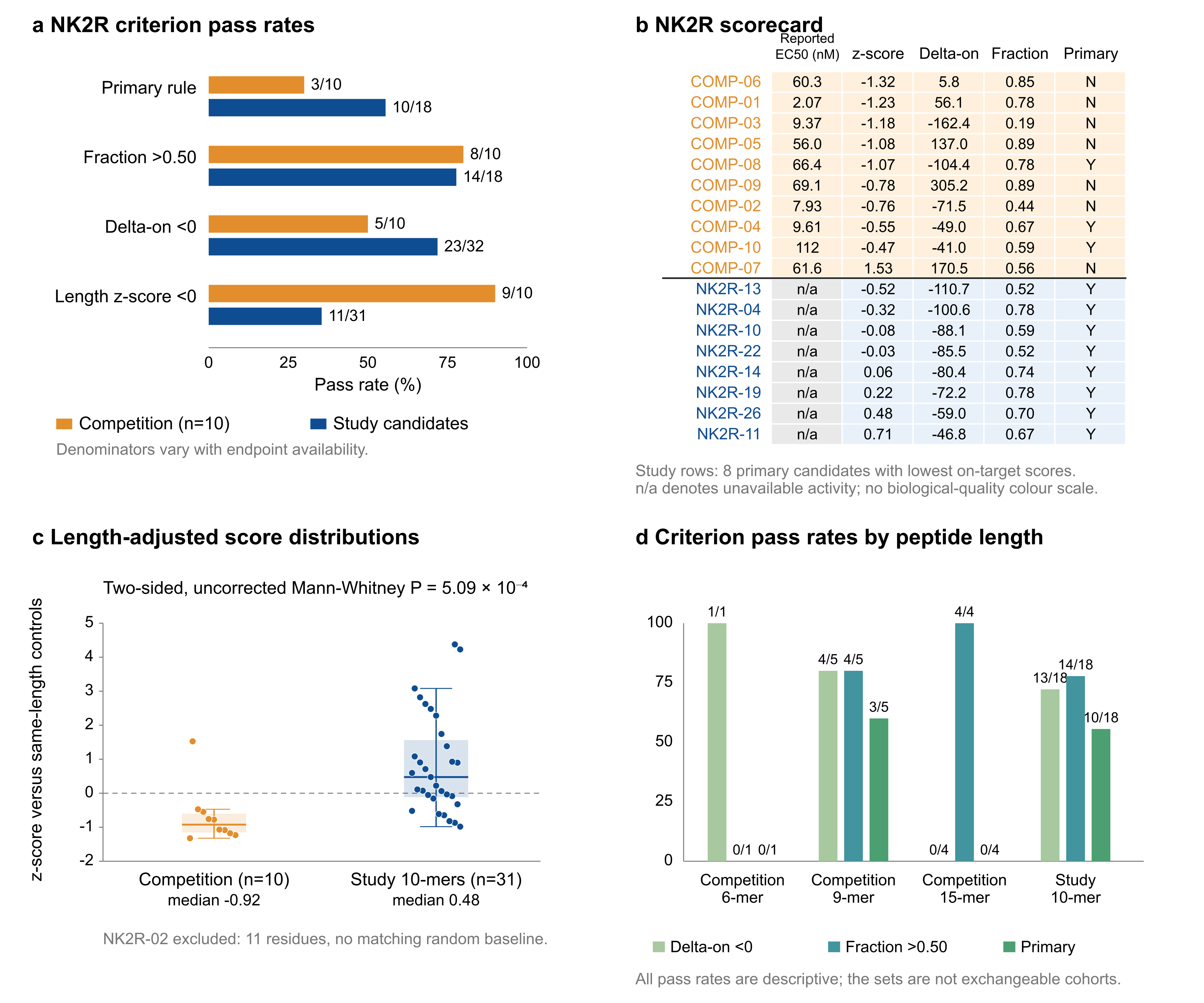

**Supplementary Fig. S7** | Descriptive NK2R comparison summaries. a, Pass counts with endpoint-specific denominators: competition n = 10; study n = 32 for Delta-on, n = 18 for neighbour-score fractions and the joint rule, and n = 31 for standardized scores. b, Scorecard for ten competition peptides and the eight primary-rule study candidates with the lowest on-target scores; n/a denotes unavailable activity. c, Standardized scores for ten competition peptides and 31 study ten-mers, excluding NK2R-02 because no eleven-residue baseline was available (competition-first U = 40, rank-biserial r = −0.742, P = 5.09 × 10⁻⁴). Boxes show medians and quartiles; whiskers extend to the furthest observations within 1.5 interquartile ranges, and all observations are shown. d, Pass counts by length. Primary denotes Delta-on < 0 and fraction > 0.50. Tests are exploratory, two-sided and uncorrected for multiplicity.

### Supplementary Fig. S8 | Extended docking-score diagnostics

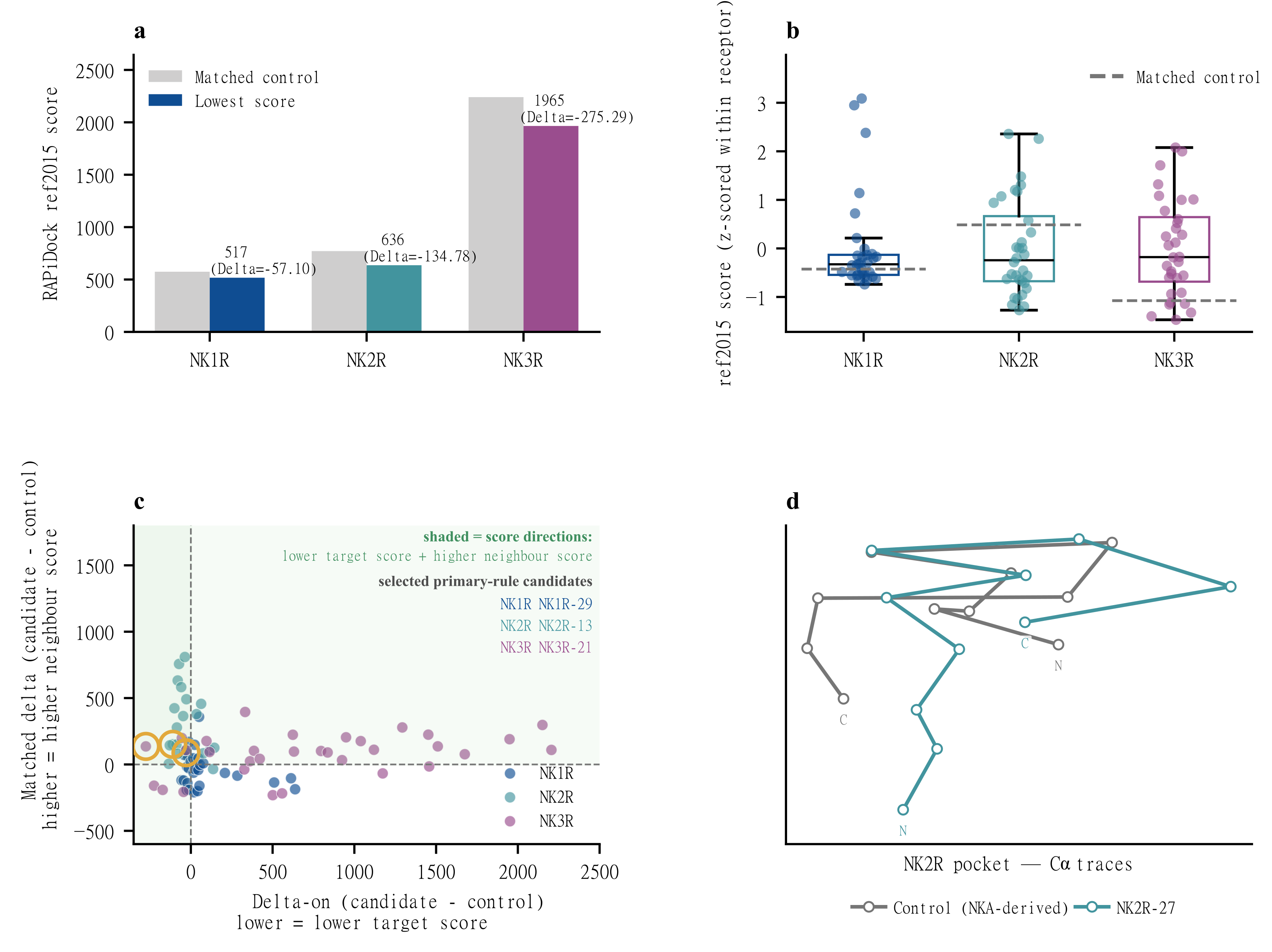

**Supplementary Fig. S8** | Extended docking-score summaries. a–b, Receptor-matched control and lowest candidate scores, candidate-minus-control differences, and within-receptor standardized score distributions. Standardization uses the mean and sample standard deviation of the 32 candidates within each receptor. c–d, Joint Delta-on and paired structural-neighbour score-difference landscape, with an archived NK2R peptide trace overlay. Shading marks the direction of the computational selection criteria.

### Supplementary Fig. S9 | Extended sequence-frequency summaries

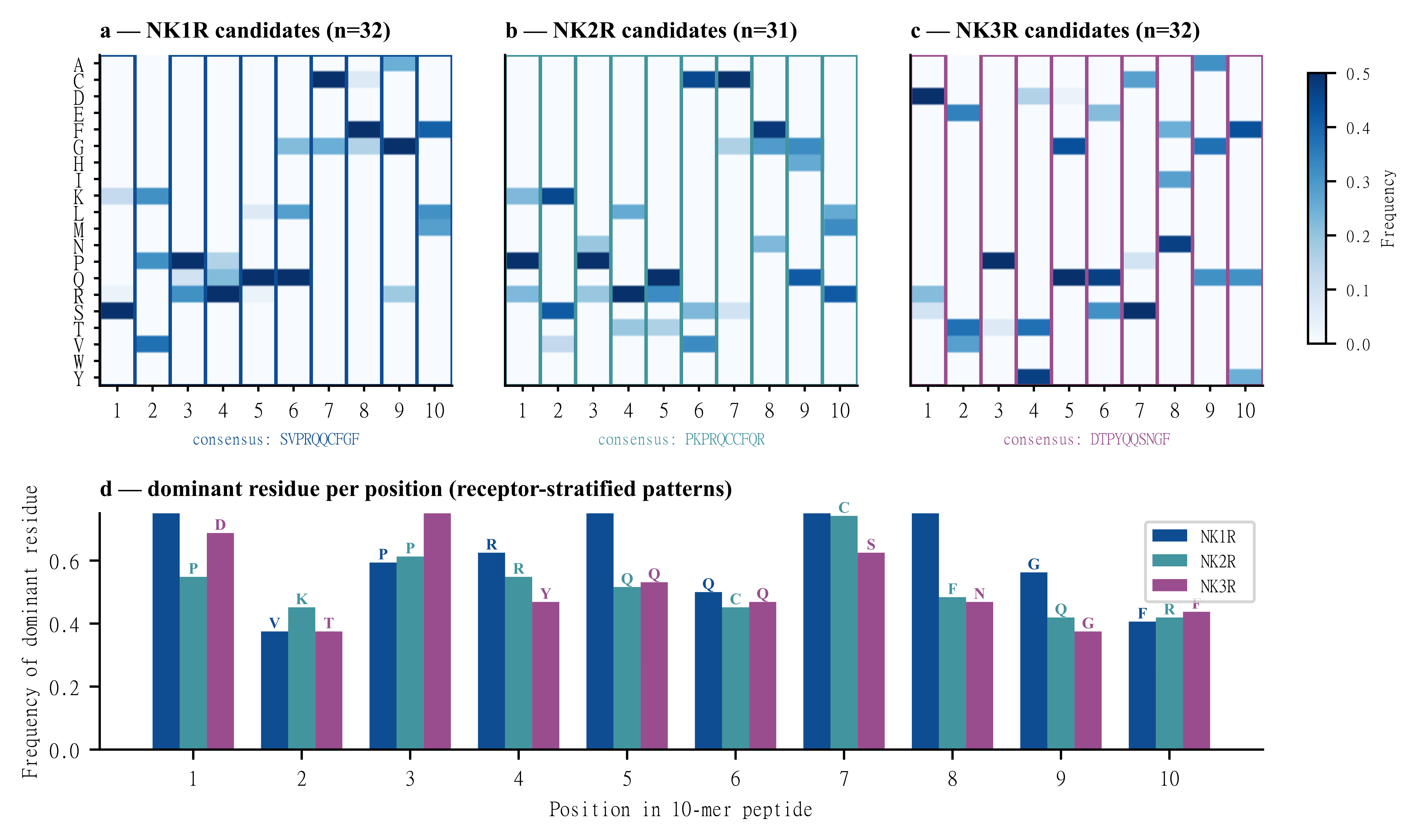

**Supplementary Fig. S9** | Extended sequence-frequency summaries. a–c, Position-wise amino-acid frequencies for exact ten-residue NK1R, NK2R and NK3R candidates; the eleven-residue NK2R-02 was excluded. d, Most frequent residue at each aligned position, stratified by receptor.
